# Cancer associated fibroblast subtypes modulate the tumor-immune microenvironment and are associated with skin cancer malignancy

**DOI:** 10.1101/2023.05.03.539213

**Authors:** Agnes Forsthuber, Bertram Aschenbrenner, Ana Korosec, Tina Jacob, Karl Annusver, Natalia Krajic, Daria Kholodniuk, Sophie Frech, Shaohua Zhu, Kim Purkhauser, Katharina Lipp, Franziska Werner, Vy Nguyen, Johannes Griss, Wolfgang Bauer, Ana Soler Cardona, Benedikt Weber, Wolfgang Weninger, Bernhard Gesslbauer, Clement Staud, Jakob Nedomansky, Christine Radtke, Stephan N. Wagner, Peter Petzelbauer, Maria Kasper, Beate M. Lichtenberger

## Abstract

Cancer-associated fibroblasts (CAFs) play a key role in cancer progression and treatment outcome. This study dissects the yet unresolved intra-tumoral variety of CAFs in three skin cancer types — Basal Cell Carcinoma, Squamous Cell Carcinoma, and Melanoma — at molecular and spatial single-cell resolution. By integral analysis of the fibroblasts with the tumor microenvironment, including epithelial, mesenchymal, and immune cells, we characterize three distinct CAF subtypes: myofibroblast-like RGS5+ CAFs, matrix CAFs (mCAFs), and immunomodulatory CAFs (iCAFs). Notably, large cohort tissue analysis reveals marked shifts in CAF subtype patterns with increasing malignancy. Two CAF types exhibit immunomodulatory capabilities via distinct mechanisms. mCAFs synthesize extracellular matrix and have the ability to ensheath tumor nests, potentially limiting T cell invasion in low-grade tumors. In contrast, iCAFs are enriched in late-stage tumors, especially infiltrative BCC and high-grade melanoma, and express unexpectedly high mRNA and protein levels of cytokines and chemokines, pointing to their integral role in immune cell recruitment and activation. This finding is further supported by our observation that in vitro exposure of primary healthy fibroblasts to skin cancer cell secretomes induces an iCAF-like phenotype with immunomodulatory functions. Thus, targeting CAF variants, particularly the immunomodulatory iCAF subtype, holds promise for improved efficacy of immunotherapy in skin cancers.

## Introduction

Fibroblasts have been considered as rather simple structural cells, which largely contribute to extracellular matrix deposition in connective tissues and tissue repair, for a long time. However, they present with an unprecedented plasticity and functional heterogeneity especially in pathological conditions, and exert a significant influence on different cells in their microenvironment, thus, modulating different processes. In cancer, fibroblasts have been established as a key component of the tumor microenvironment (TME) affecting both cancer progression and the response to therapies. Fibroblast heterogeneity has been acknowledged previously as playing both tumor suppressive as well as tumor supportive roles ^1–7^. Recent studies even demonstrated that also mutations in fibroblasts can lead to cancer^8^, further highlighting their impact on tumorigenesis. It also has become clear that in a single tumor several fibroblast subtypes exist in parallel with different functions ^9^. Single-cell RNA sequencing (scRNA-seq) has revealed manifold dermal fibroblast subpopulations in mouse and human healthy skin ^10–12^. Likewise, fibroblast heterogeneity has been studied in many cancers such as breast cancer ^13,14^, pancreatic ductal adeno carcinoma ^15,16^, colorectal cancer ^17,18^, head and neck squamous cell carcinoma ^19^ and many more ^20^. Comparable single-cell transcriptomics studies in skin cancer are missing. The few previous scRNA-seq based studies of human melanoma, basal cell carcinoma (BCC) and cutaneous squamous cell carcinoma (SCC) mainly focused on tumor infiltrating lymphocytes, thus these datasets included only a few or no fibroblasts (Supplemental Table 1) ^21–27^. In the past years, scRNA-seq revealed a range of different cancer-associated fibroblast (CAF) subsets with newly described markers in various tissues. Besides context-dependent and/or uniquely described CAF subsets in certain types of cancer ^28–30^, a common trait among many of these studies is the presence of a CAF subtype with immunomodulating characteristics, secreting IL6 and other proinflammatory cytokines, and a CAF subtype with a myofibroblastic phenotype defined by its widely accepted signature molecule alpha smooth muscle actin (ACTA2) ^13,14,31–36^. In healthy skin, lineage tracing mouse models and functional studies in mouse and human skin identified two major fibroblast subsets, the papillary fibroblasts in the upper dermis and the reticular fibroblasts of the lower dermis. They exhibit distinct roles during skin development, wound healing and fibrotic skin disorders, ^37–40^. Whether these two subpopulations evolve into CAFs, or impact skin tumor progression differently, is still an unresolved topic. So far, all studies on skin CAFs focused on CAFs as a unit, their overall marker gene expression, capability to stimulate migration of tumor cells, secretion of soluble mediators, and their response to anti-tumor therapies ^41–43^, leaving a major gap in knowledge about the presence of CAF subtypes, their association with tumor malignancy and different functional roles, which we address in this study.

Here, we investigated the cellular ecosystem of BCC, SCC and melanoma – with emphasis on fibroblast heterogeneity – using the sensitive Smart-seq2 scRNA-seq platform (n=10 tumors) and mRNA staining *in situ* (n=68 tumors). SCCs vary from well to poorly differentiated with increasing metastatic potential ^44,45^, and BCCs, which metastasize very rarely, are primarily tissue-destructive locally ^46,47^. Both cancer types arise from keratinocytes. In contrast, melanoma is a skin cancer type derived from melanocytes that comes with a high potential of metastatic spread and poor survival rates, which have improved significantly due to novel therapeutics, especially immunotherapies ^48,49^. The unique combination of these three different types of skin tumors in one dataset allowed us to dissect similarities and differences between those cancer types with a special focus on the fibroblast heterogeneity in a healthy skin, premalignant and cancer context. Collectively, our work deconstructed CAF heterogeneity in skin cancer, which led to important new insights. We show that the mCAF subtype, which forms a dense extracellular matrix network at the tumor-stroma border, likely plays a role in T cell marginalization, and propose that the iCAF subtype is an important regulator of immune cell recruitment and immune surveillance.

## Results

### A collective single-cell atlas of primary BCC, SCC and melanoma to deconstruct skin cancer

To explore fibroblast heterogeneity and their cross talk to tumor and immune cells, we collected biopsies from four cutaneous squamous cell carcinomas (SCCs), three basal cell carcinomas (BCCs), three melanomas and biopsies of sun-protected skin from five healthy donors (Supplemental Table 2). For the tumor samples, fresh 4 mm punch biopsies from the tumor center and from non-lesional adjacent skin (providing sex- and age-matched healthy skin controls) were collected directly after surgery. Since previous single cell transcriptomic studies of skin cancer included no fibroblasts or only low CAF numbers (Supplemental Table 1), instead of random droplet-based sampling of tumor-associated cells, we chose a FACS-sorting approach to enrich for fibroblasts and to gain highly sensitive scRNA-seq data using Smart-seq2 technology. Upon dissociation of the tissues, the samples were enriched for keratinocytes, fibroblasts and immune cells by FACS (Figure S1A and S1B) and the cells were directly sorted into 384-well plates for sequencing with the Smart-seq2 technology (Figure 1A). In two healthy control samples a random live-cell sorting approach was used, explaining why keratinocytes are underrepresented in these samples (Healthy I, Healthy II, Figure 2A), which was compensated by a separate keratinocyte enrichment protocol in healthy samples Healthy III-V (Figure S1A and 2A). In total, 5760 cells were sequenced, finally retaining 4824 cells at a median depth of 486146 RPKM/cell and a median of 3242 genes/cell after quality control and filtering (Figure S1C). Cell numbers per sample after quality filtering are shown in Figure S1D. Unsupervised clustering separated cells into fibroblasts, healthy and malignant keratinocytes as well as melanocytes, immune cells, and endothelial cells (Figure 1B). Assignment of cell type identity was based on commonly accepted signature gene expression: *COL1A1* and *PDGFRA* for fibroblasts, *RERGL* for vascular smooth muscle cells (vSMC), *KRT5*, *KRT14* and *CDH1 (E-cadherin)* for keratinocytes, *CD45* for immune cells, *TYR*, *MITF* and *MLANA* for melanocytic cells and *CDH5* for endothelial cells (Figure 1B and 1C).

**Figure 1.**
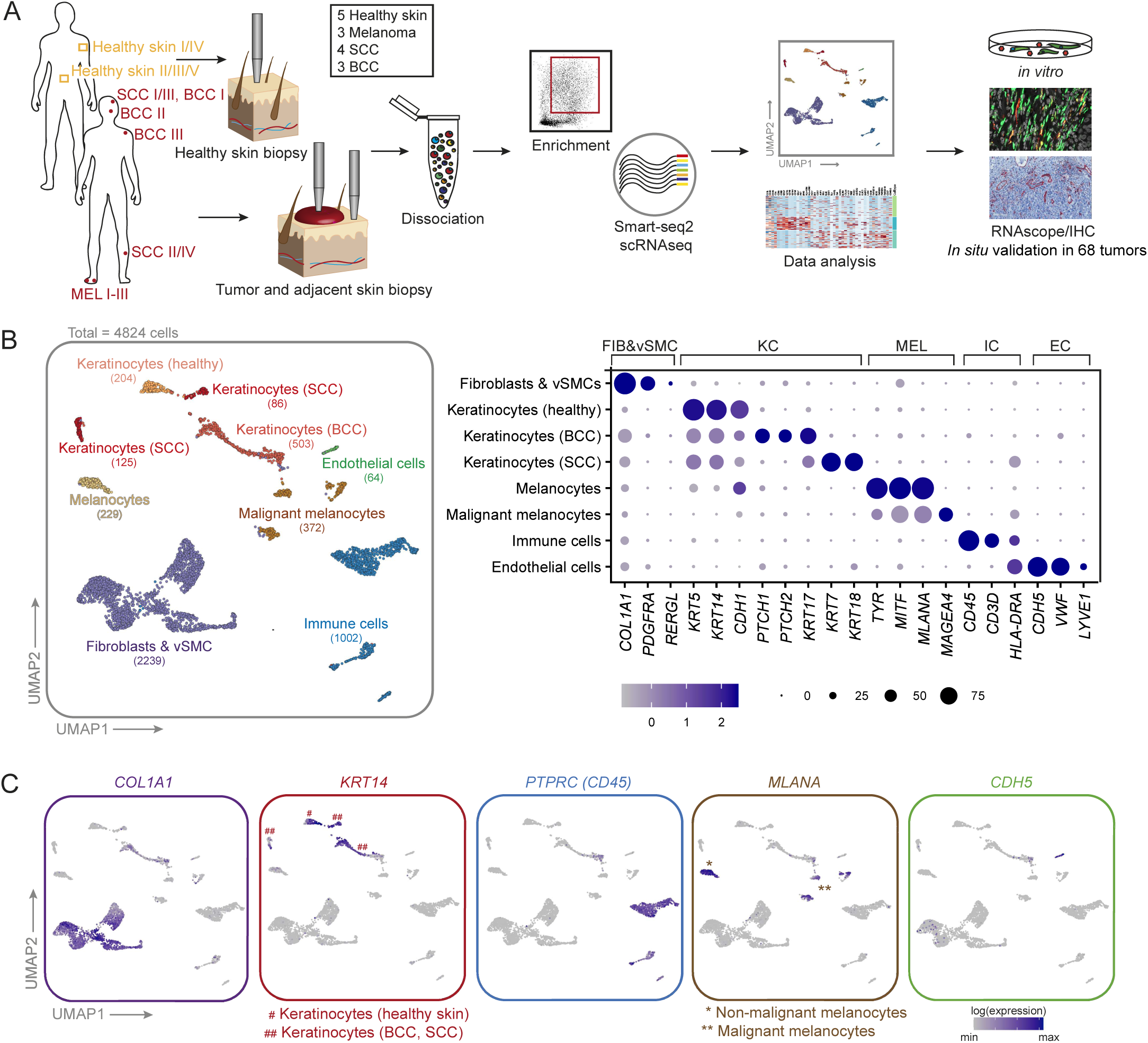
A single cell transcriptomic atlas of human BCC, SCC, melanoma and healthy skin. (A) Workflow of donor sample processing for Smart-seq2 scRNA-seq, data analysis and verification. (B) UMAP projection of first-level clustering of 4824 cells (left). Clusters are labelled by cell types, which were identified by commonly accepted marker genes (right). (C) Expression of top marker genes for the main cell types.

**Figure 2.**
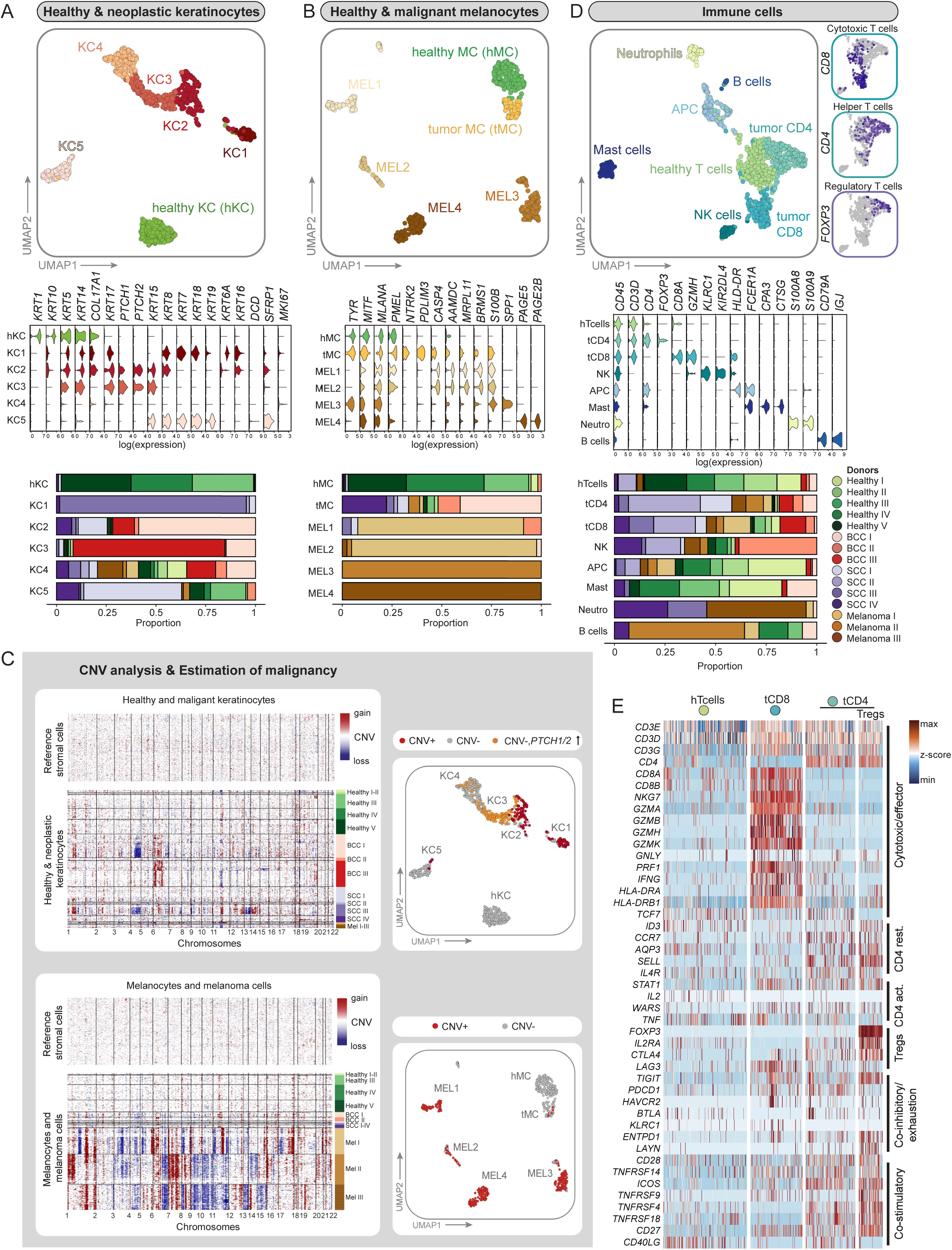
Second-level clustering of non-mesenchymal cells and CNV analysis. (A-B) UMAP projection of second-level clustering, violin plots of signature genes as well as bar plots showing donor sample distribution per cluster are presented for healthy and neoplastic keratinocytes and melanocytes. (C) CNV analysis (based on *inferCNV* package) of tumor samples using stromal cells as reference controls. UMAPs for healthy and neoplastic keratinocytes and melanocytes: Malignant cells with a predicted CNV alteration are highlighted in red and *PTCH1*/*PTCH2* overexpressing cells without CNVs are highlighted in orange or yellow, respectively. (D) UMAP projection of second-level clustering, violin plots of signature genes as well as bar plots showing donor sample distribution per cluster are presented for immune cells. The distribution of cytotoxic, helper and regulatory T cells are depicted in separate cut-outs. (E) Heatmap of genes that reflect the resting, activation, cytotoxic, co-stimulatory or co-inhibitors status of T cell subsets from healthy and tumor samples. KC-keratinocyte, MC-melanocyte, MEL-melanoma cells, hTcells-healthy T cells, tCD8-Cytotoxic T cells, tCD4-Helper T cells, Tregs-Regulatory T cells.

### Tumor cells express a range of additional keratins

Second-level clustering of healthy and neoplastic keratinocytes resulted in a clearly separated cluster of healthy keratinocytes (hKC) with basal (*KRT5*, *KRT14*, *COL17A1*) and differentiating/suprabasal (*KRT1*, *KRT10*) marker gene expression. The majority of cells in clusters KC1–KC5 contained tumor cells from BCC and SCC samples, with tumor cells expressing a range of additional keratins that are not expressed by healthy skin keratinocytes (Figure 2A and Supplementary table 2). For example, BCC cells expressed *KRT5*, *KRT14* and additionally *KRT17* ^50^, which are foremost clustering in KC2 and KC3. As expected for the Hedgehog(Hh)-pathway-dependent BCC, we confirmed *PTCH1/2* as well as *GLI1/2* overexpression in clusters KC2 and KC3 (Figure 2A, S2A and S2B) ^51^. Cells from the SCC samples clustered in KC1, KC2, KC4 and KC5. KC1 – which primarily comprised cells of donor sample SCC III (Figure 2A and S2C) – expressed in addition to *KRT5*, *KRT14*, and *KRT17* the typical SCC keratins like *KRT6A* and *KRT16,* and showed reduced expression of *KRT1* and *KRT10* ^50^ (Figure 2A). Cells from donor sample SCC IV, a SCC arisen from Bowen’s disease (Supplemental Table 2), mainly clustered in KC2, KC4, and KC5 that showed reduced expression of *KRT1*, *KRT10*. KC2 and KC5 also displayed expression of *KRT16* and *KRT19,* respectively, which has been previously described for this SCC subtype ^50^ (Figure 2A and S2C). Notably, *KRT7*, *KRT18* and *KRT19* were only expressed by KC1 and KC5, which were primarily represented by cells originating from SCC samples (Figure 2A and S2C). It has been shown that tumors co-opt developmental programs for their progression ^1^. In our dataset we find previously described keratins of the simple skin epithelium of the first trimester embryo such as KRT18 and KRT19 expressed in SCC cluster KC1 and KC5, or *KRT8* expressed in BCC cluster KC2 ^52^ (Figure 2A).

Interestingly, also healthy keratinocytes contributed with more than 25% to KC5, which likely represent luminal cells from sebaceous or sweat glands expressing the marker genes *KRT7*, *KRT19*, *SFRP1* and *DCD* ^53^. In KC4, keratinocytes from BCC, SCC, melanoma as well as healthy skin clustered together. These cells did not express unified classical keratinocyte markers (Figure 2A) but presented with a scattered expression of different hair follicle-associated keratins (data not shown). This expression pattern suggests that KC4 contains keratinocytes of the different anatomical structures of the hair follicle in line with KC4 cluster’s position next to the BCC keratinocytes (KC3) and the presence of *PTCH1/2* and *GLI1/2* expressing cells (Figure 2C and Figure S2 A-B).

### The tumor microenvironment strongly impacts non-malignant cell types

Second-level clustering of melanocytes and melanoma cells revealed one cluster of healthy melanocytes (hMC) derived from healthy skin samples, and one separate cluster of non-malignant melanocytes derived mainly from SCC and BCC samples (tumor melanocytes, tMC). This separate clustering can likely be explained by the influence of the TME on the expression pattern of melanocytes in comparison to melanocytes from healthy skin. The melanoma samples separated into four different donor-specific clusters: MEL1 and MEL2 coming from Melanoma I, MEL3 from Melanoma II, and MEL4 from Melanoma III (Figure 2B and Supplementary table 2). In order to explain this donor-specific clustering, as well as to segregate malignant from non-malignant cells, we inferred copy number variation (CNV) analysis on our scRNA-seq data using the R package *inferCNV* ^19,25^. We performed CNV analysis for melanocytes and melanoma cells, and a separate CNV analysis for healthy and malignant keratinocytes, as previously described ^19^, using healthy stromal cells (fibroblasts, vascular smooth muscle cells and pericytes) as a reference (Figure 2C and S3). Melanoma cells, which are known for their high mutational load ^54^, displayed strong CNV patterns. Remarkably, these patterns were donor-specific despite of sharing the same histological subtype (acral lentiginous melanoma, ALM) and body location (heel/toe) (Supplementary table 1). Among BCC and SCC samples, SCC III (forming cluster KC1) presented with the strongest CNV pattern, which matched well with its histopathological characterization of a poorly differentiated, aggressive type of SCC (Supplementary table 2). In addition to the CNV analysis, we also utilized high *PTCH1* and *PTCH2* expression^55,56^ (*PTCH1/2 high)* in comparison to the healthy skin cluster hKC to pinpoint tumor cells (Figure 2A and S2A,B). As the non-malignant cells mixed across almost all donor samples, such as tMC or immune cells (Figure 2B-D), we did not regress for donor differences; thus, donor-specific tumor cell clustering in KC1 and MEL1–MEL4 is likely the result of genomic aberrations and not an effect of batch variations.

Subclustering of immune cells resulted in eight immune cell clusters, including T and B cells, granulocytes and antigen presenting cells (Figure 2D and S2C). Interestingly, CD4 and CD8 T cells from healthy samples (hTcells) formed a distinct cluster, adjacent to CD4 (tCD4) and CD8 T cells (tCD8) from tumor samples (Figure 2D and S2C). This can be explained by a different activation status of T cells in healthy or tumor samples: Increased expression of granzymes, perforin and *IFNγ* indicated T cell activation in CD8 T cells from tumor samples versus healthy tissue (Figure 2E). Regulatory T cells (Tregs), *CD4*^+^*IL2RA*^+^*FOXP3*^+^, were found mixed with CD4 T cells from tumor samples (tCD4) (Figure 2D, 2E and S2C).

### Fibroblasts from healthy skin cluster separately from CAFs

Second-level clustering of fibroblasts and vSMCs (2239 cells) resulted in 7 subclusters: two main clusters from healthy skin samples (papillary fibroblasts (pFIB), reticular fibroblasts (rFIB)), four CAF subclusters (matrix CAFs (mCAF), immunomodulatory CAFs (iCAF), *RGS5^+^* CAFs and pericytes (*RGS5^+^*cells), unclassifiable CAFs (ucCAF)), and one vSMC cluster (Figure 3A, S4A and S4B). We used differentially expressed genes as well as commonly accepted markers to define these subclusters. Reassuringly, fibroblasts from the tumor-adjacent skin samples were found on the transition between healthy and tumor samples, and within the *RGS5^+^*as well as vSMC cluster (Figure S4A), indicating that CAFs may develop from skin-resident fibroblasts (pointing towards field cancerization) ^57^.

**Figure 3.**
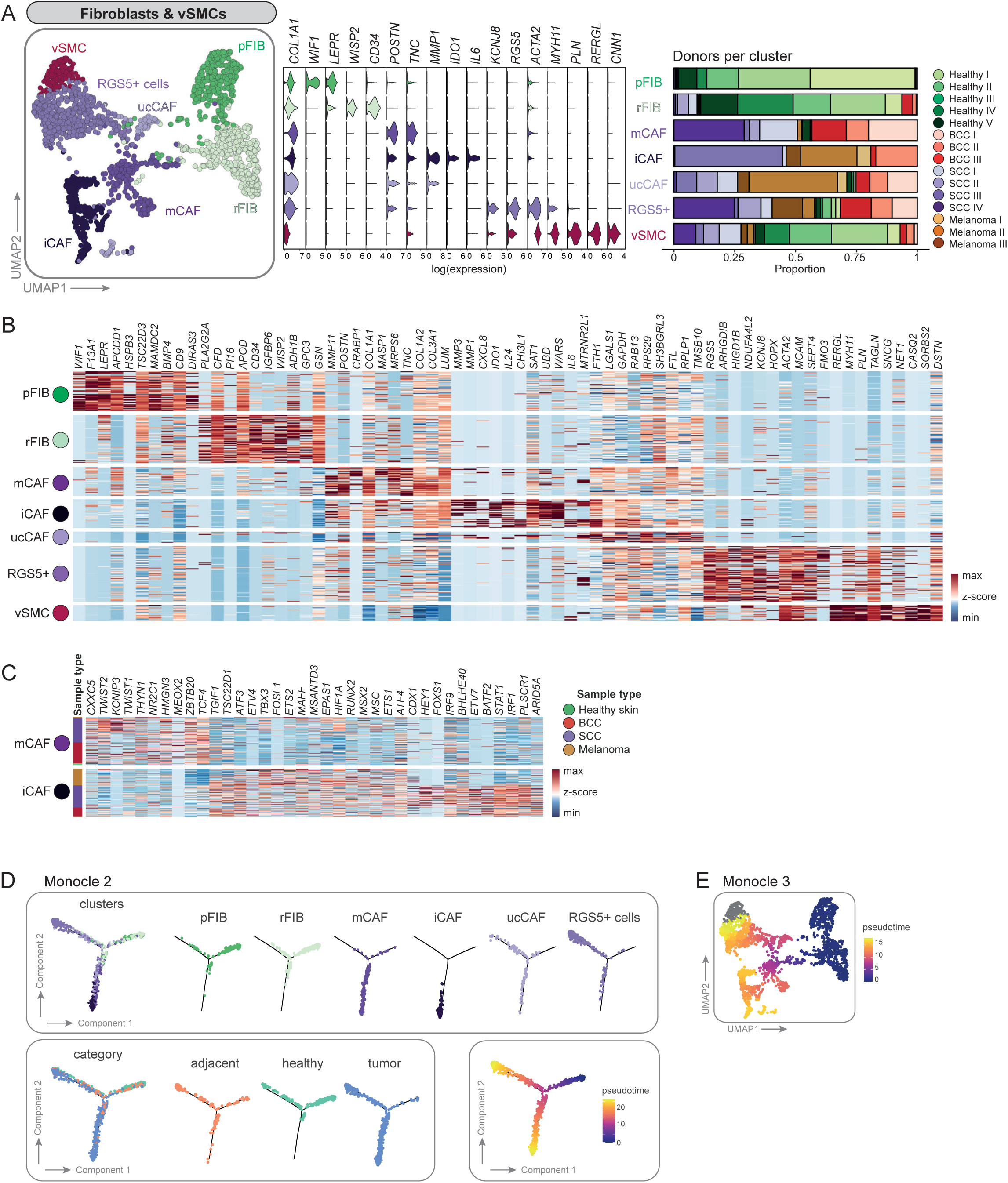
Second-level clustering of fibroblasts and vascular smooth muscle cells (vSMCs) results in two heathy fibroblasts populations, four CAF subsets and one vSMC cluster. (A) UMAP of second-level clustered fibroblasts and vSMCs. Violin plots of signature genes and bar plots showing donor sample distribution per cluster. (B) Heatmap of top ten differentially expressed genes per cluster. (C) Differentially expressed transcription factors between mCAFs and iCAFs. (D) Trajectoy analysis using Monocle2. Cells were highlighted according to clusters, category or pseudotime. (E) UMAP colored in pseudotime showing trajectory results from Monocle3

Importantly, several established markers for papillary and reticular fibroblasts could be assigned to the two subclusters comprising cells from healthy donors. *COL6A5*, *COL23A1* and *HSPB3* ^10^, the Wnt inhibitors *APCDD1* and *WIF1* ^10,58^, *NTN1* and *PDPN* ^59^ are commonly accepted markers for papillary fibroblasts and were found to be expressed in the pFIB cluster. *DPP4* (*CD26*), which was described as a marker for papillary fibroblasts by Tabib et al. 2018, was expressed in both subpopulations in our data (Figure S4C), as shown by Korosec et al., 2019 and Vorstandlechner et al., 2020 ^12,39^. The healthy fibroblast subcluster rFIB expressed genes that are characteristic for reticular fibroblasts: *THY1* ^39^, *FMO1*, *MYOC* and *LSP1* ^11^, *MGP* and *ACTA2* ^59^ as well as the preadipocyte marker genes *PPARG* and *CD36* ^39^ (Figure S3C). However, we could not confirm the expression of various other published reticular fibroblast markers. The discrepancy of marker expression in distinct datasets might result from tissue collection from different body sites, tissue preparation or sequencing technology. Furthermore, several markers were identified in *in vitro* cultures, mostly on protein level. Thus, we conclude that the healthy fibroblast cluster pFIB represents papillary fibroblasts, and rFIB represents reticular fibroblasts (Figure 3A, 3B and S4C).

### Different skin cancer types comprise both common and tumor type-specific CAF subsets

Subclustering segregated four CAF populations: mCAFs, iCAFs, *RGS5^+^* cells, and ucCAFs. Matrix CAFs (mCAFs) exhibited increased expression of extracellular matrix components such as collagens (*COL1A1*, *COL1A2*, *COL3A1)*, Lumican (*LUM*), Periostin (*POSTN*) or Tenascin-C (*TNC*) compared to all other fibroblast clusters (Figure 3A and 3B). Immunomodulatory CAFs (iCAFs) presented with enhanced expression of the matrix remodelers *MMP1* and *MMP3*, the pro-inflammatory cytokines *IL6* and *CXCL8* and the immune-suppressive molecule *IDO1* among their top ten differentially expressed genes (DEGs), thus suggesting an immunoregulatory and cancer invasion-supportive phenotype (Figure 3A and 3B). Intriguingly, the mCAF cluster harbored fibroblasts from all BCC and well-differentiated SCC samples (mainly SCC I and SCC IV), whereas the iCAF cluster contained fibroblasts from all melanoma samples, one poorly differentiated SCC (SCC III) and one BCC (BCC II). While the presence of these subsets seems to depend on the skin cancer type with iCAFs being associated with the most aggressive tumors, *RGS5^+^* cells were found in all tumor samples independently of skin cancer type and malignancy (Figure 3A, S4A and S4B). Notably, mCAFs and iCAFs expressed different transcription factors (TFs; Figure 3C). In mCAFs, TFs associated with conserved developmental proteins, including genes of the WNT pathway (*CXXC5*, *TCF4*), transcriptional regulation of mesenchymal cell lineages (*TWIST1*, *TWIST2*), and anti-inflammatory signaling (*KCNIP3*), were upregulated. iCAFs expressed high levels of TFs that are related to immune responses such as *STAT1*, *IRF1*, *IRF9* or *ARID5A*. Of note, there is also a difference in TF expression between SCC- and melanoma-derived iCAFs.

Since *RGS5^+^* cells express *ACTA2* (in combination with *COL1A1*, Figure 4A), this subset likely represents activated fibroblasts that are usually termed myofibroblasts in wounded or fibrotic tissues ^60^, or myoCAFs in different cancer types ^35^. However, they also express genes among their top DEGs that have been used as pericyte markers, such as *RGS5*, *KCNJ8, ACTA2* and *MCAM* (Figure 3B) ^61–63^. *RGS5^+^*cells also share some markers with the vSMC cluster, such as *ACTA2*, *TAGLN* and *MCAM*, which are markers for perivascular cells in various tissues ^39,64^. The vSMC population formed its separate cluster and was clearly defined by *RERGL*, *MYH11*, *CNN1* in addition to *ACTA2* ^62^ (Figure 3A and 3B). Unclassifiable CAFs (ucCAFs) represent a minor population with an inconclusive gene expression pattern and mixed cell contribution from almost all tumor samples, which we therefore did not consider further for in-depth discussion (Figure 3A and 3B).

**Figure 4.**
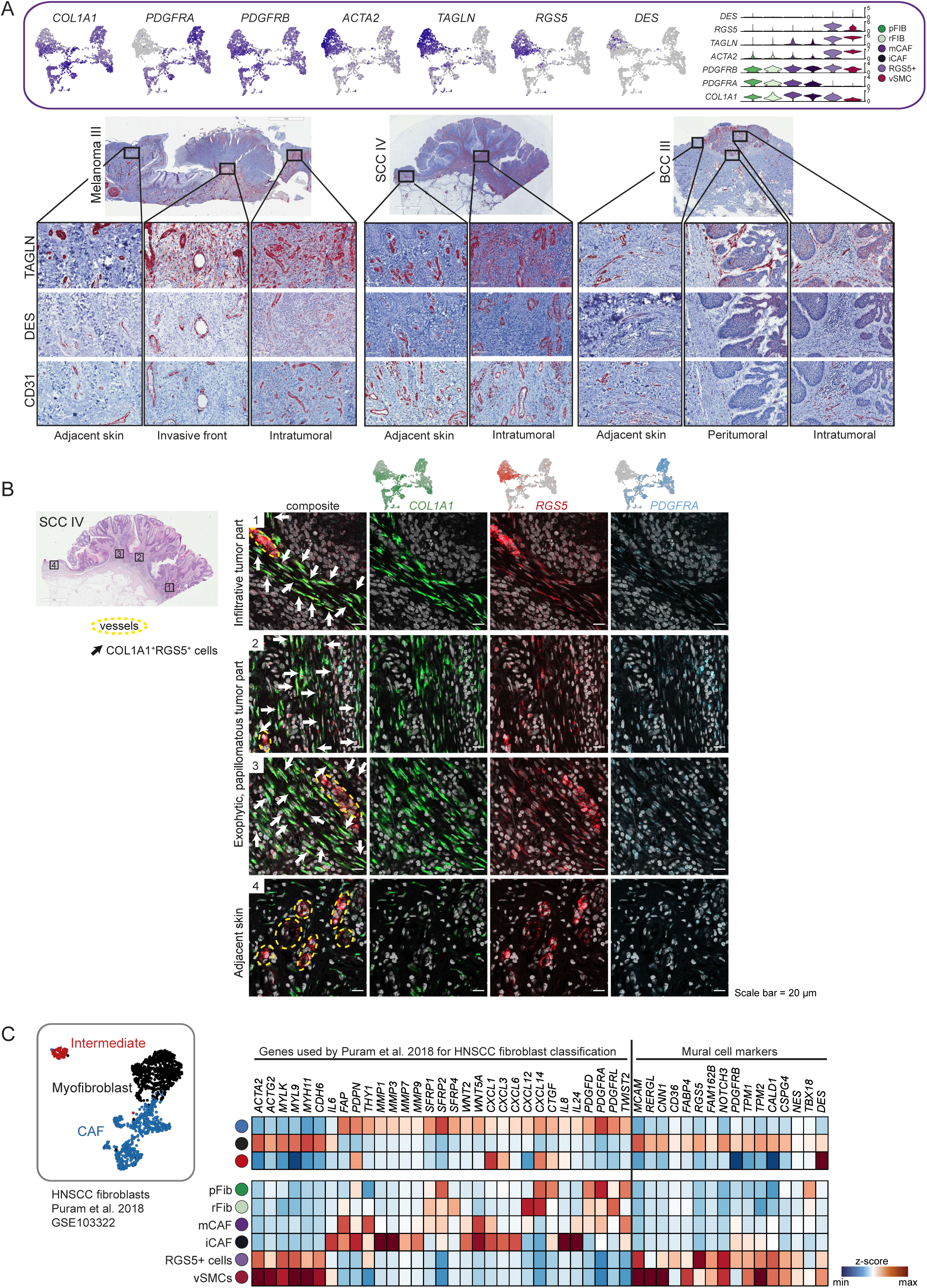
The RGS5^+^ cells are an inhomogeneous population of CAFs and pericytes. (A) Feature and violin plots showing the expression of fibroblast and pericyte marker genes in the RGS5^+^ cluster. Representative immunohistochemistry of TAGLN, DES and CD31 in different regions of the tumor (intratumoral, peritumoral). (B) Representative images of *COL1A1* (green), *RGS5* (red) and *PDGFRA* (blue) RNAScope fluorescence stainings in four different regions of FFPE tissue sections from donor sample SCC IV. DAPI nuclear stain is shown in grey. Scale bar represents 20 μm. (C) Myofibroblasts in a HNSCC dataset from Puram et al. 2017, exhibits a very similar expression pattern in comparison to the RGS5^+^ cluster in our dataset.

### Trajectory inference shows two main differentiation routes for CAFs from healthy cells

Trajectory analysis using Monocle2 and Monocle3 ^65,66^ showed that healthy fibroblasts follow two differentiation routes: either towards mCAF/iCAF or towards *RGS5*^+^cells (Figure 3D and 3E). Note that the vSMC cluster was excluded from trajectory inference as we do not expect them to differentiate from healthy fibroblasts and CAFs from a biological point of view. The trajectory analysis also shows that iCAFs are a differentiation endpoint, with mCAFs being an intermediate state, and thus it may be possible that iCAFs develop from mCAFs. This might also be supported by the fact that iCAFs express several mCAF genes albeit at a lower level, while mCAFs do not express iCAF markers (Figure S8A and S8B). Fibroblasts from tumor adjacent skin samples are preferentially found in the healthy fibroblast and the RGS5^+^ cells branches, and a smaller fraction bridging to the mCAF branch, indicating that they are in a transitory position between healthy fibroblasts and CAFs.

### *ACTA2* and *FAP* in combination identify all CAF subpopulations in skin tumor samples

Previous studies have used different single markers like ACTA2 or FAP to identify or isolate CAFs in different tissues ^67^. However, a query on previously described CAF marker genes showed that, in our dataset, most of them are either restricted to a distinct CAF subset, or found in all fibroblast clusters including healthy fibroblasts, but do not solely identify all CAF subsets (Figure S4D). A good strategy to detect all CAFs within skin tumor samples is combining the two most frequently used CAF markers *ACTA2* and *FAP*. Although this combination also includes vSMCs, it allows to enrich for all CAF subpopulations when used together (Figure S4D).

### The *RGS5^+^* cells are best described as a mixed population of myoCAFs and pericytes

When we investigated the detailed gene expression profiles of the distinct CAF subtypes we suspected that the *RGS5^+^* cluster likely comprises both myoCAFs and pericytes, therefore we chose the neutral term “*RGS5^+^* cells” for this cluster. In detail, *RGS5^+^* cells expressed *ACTA2 –* the signature gene for myofibroblasts and myoCAFs – in combination with *COL1A1*. The expression of *PDGFRB*, *TAGLN*, *RGS5*, *DES* and the absence of *PDGFRA* suggests that this cluster also comprises pericytes ^61–63^ (Figure 4A). Interestingly, the *RGS5^+^* cluster also shows expression of *NOTCH3*, *EPAS1*, *COL18A1* and *NR2F2*, markers that were used to describe so-called vascular CAFs (vCAFs), a CAF subset defined by Bartoschek et al., 2018 in a mouse model for breast cancer ^68^ (Figure S5A). Of note, endothelial markers such as *CDH5*, *PECAM1*, *TIE1* or *CD62* were not expressed in the *RGS5^+^* cluster (Figure S5B).

To characterize the nature of the *RGS5^+^* cluster further, we stained tumor sections for Transgelin (TAGLN), which is a prominently expressed gene in this cluster. This tissue staining revealed TAGLN protein along blood vessels as expected, but also within the tumor stroma without direct contact to vessel-like structures (Figure 4A). Protein expression of Desmin (DES), a marker for pericytes and vSMCs ^60,61^ was restricted to vessels (Figure 4A). *DES* was only expressed by few cells on RNA level, which was not sufficient to identify a separate pericyte cluster within the *RGS5^+^* cells in our sequencing data set ^69,70^. Next, we performed mRNA co-staining for *RGS5, COL1A1* and *PDGFRA* mRNA to validate our sequencing data and to verify the stromal and perivascular presence of *RGS5^+^* cells (Figure 4B, Figure S5C). In tumor regions (Figure 4B region 1-3 and S4C region 1-2), a positive staining for *RGS5* was detected both at vessel-structures and within the stroma, in comparison to the peritumoral area, where *RGS5* staining was only found surrounding vessel-like structures (Figure 4B, region 4).

To shed light on the discrepant classification of cells as being myofibroblast-like CAFs or pericytes in tumor samples, we reanalyzed a publicly available head and neck squamous cell carcinoma (HNSCC) data set ^19^ and put it in comparison with our dataset. Puram et al. classified the tumor fibroblasts into CAFs, myofibroblasts and intermediate (resting) fibroblasts. We extended their published marker gene set by commonly accepted pericyte markers and found those enriched in the myofibroblast cluster only, revealing a very similar expression profile to our *RGS5^+^* cell cluster (Figure 4C). The fact that this formerly defined myofibroblasts have been defined as pericytes upon reanalysis by another group ^71^, suggests that myofibroblasts and pericytes share a very close gene expression pattern which indeed does not allow segregation by transcriptional profiling. Thus, the absence of histological stainings in previously published datasets impeded an accurate definition of those cells, and only the combination of histological localization and gene expression allows proper lineage designation. We conclude that the *RGS5^+^* cell cluster within our as well as the HNSCC dataset comprises both pericytes and CAFs.

### *In situ* validation and spatial localization of mCAF and iCAF subsets

We verified the presence of iCAFs and mCAFs by mRNA staining *in situ* in the same tumor samples that were sequenced as well as in additional independent tumor biopsies (n=68 tumors in total). We used *COL11A1* and *PTGDS* as markers for mCAFs, and MMP1 (and several cytokines) as a marker for iCAFs, in co-stainings with the pan-fibroblast marker *COL1A1* (Figure 5A, 5B, 6C, S6 and S7). The distribution of mCAFs and iCAFs in the tumor tissue follows different patterns: mCAFs were found abundantly in large patches ensheathing tumor nests, but also pervading the tumor in strands (Figure 5A and S6A). Importantly, mCAFs were especially enriched at the tumor-stroma border of BCC and well-differentiated SCC (Figure S6A). Contrary, iCAFs were found in smaller numbers intermingled between *MMP1*^-^*COL1A1*^+^ cells intratumorally in stromal nests and strands that pervade the tumors or in patches at the invasive front (Figure 5B and S6B). To verify our scRNA-seq data suggesting that iCAFs are predominant in aggressive tumors, we stratified the tumor samples into different categories: nodular and infiltrative BCC, well-differentiated and poorly-differentiated SCC, and low-grade (Tis and ≤T1) and high-grade (≥ T3) melanomas (n=52). Large-field spatial visualization of the CAF subpopulations in tumor tissue samples showed a clear change in the CAF patterns from lower to higher malignancy, along with a higher overall CAF density in the aggressive variants of the respective skin cancer subtypes (Figure 5C). To quantify this difference in CAF subsets, the regions of interest (ROIs) were set within the tumor as well as at the invasive front (Figure 5C,D and S7A; see Methods). The total CAF density significantly increased in infiltrative BCC compared to nodular BCC, and in high-grade melanoma compared to low-grade melanoma (Figure 5D). Also the iCAFs displayed an increase in number between nodular BCC and infiltrative BCC, and low grade (≤ T1) and high grade (≥ T3) melanomas, respectively (Figure 5D). The SCC subtypes displayed a similar iCAF trend; however, it was not statistically significant (see Discussion). This extended data analysis, which also included infiltrative BCC and low-grade melanoma samples, confirmed the scRNA-seq data showing that iCAFs are more abundant in more malignant skin cancer subtypes, particularly in infiltrative BCC and high-grade melanoma. Interestingly, also the mCAFs increased in abundance in infiltrative BCC compared to nodular BCC, and high grade (≥ T3) versus low grade (≤ T1) melanomas (Figure 5D, Discussion).

**Figure 5.**
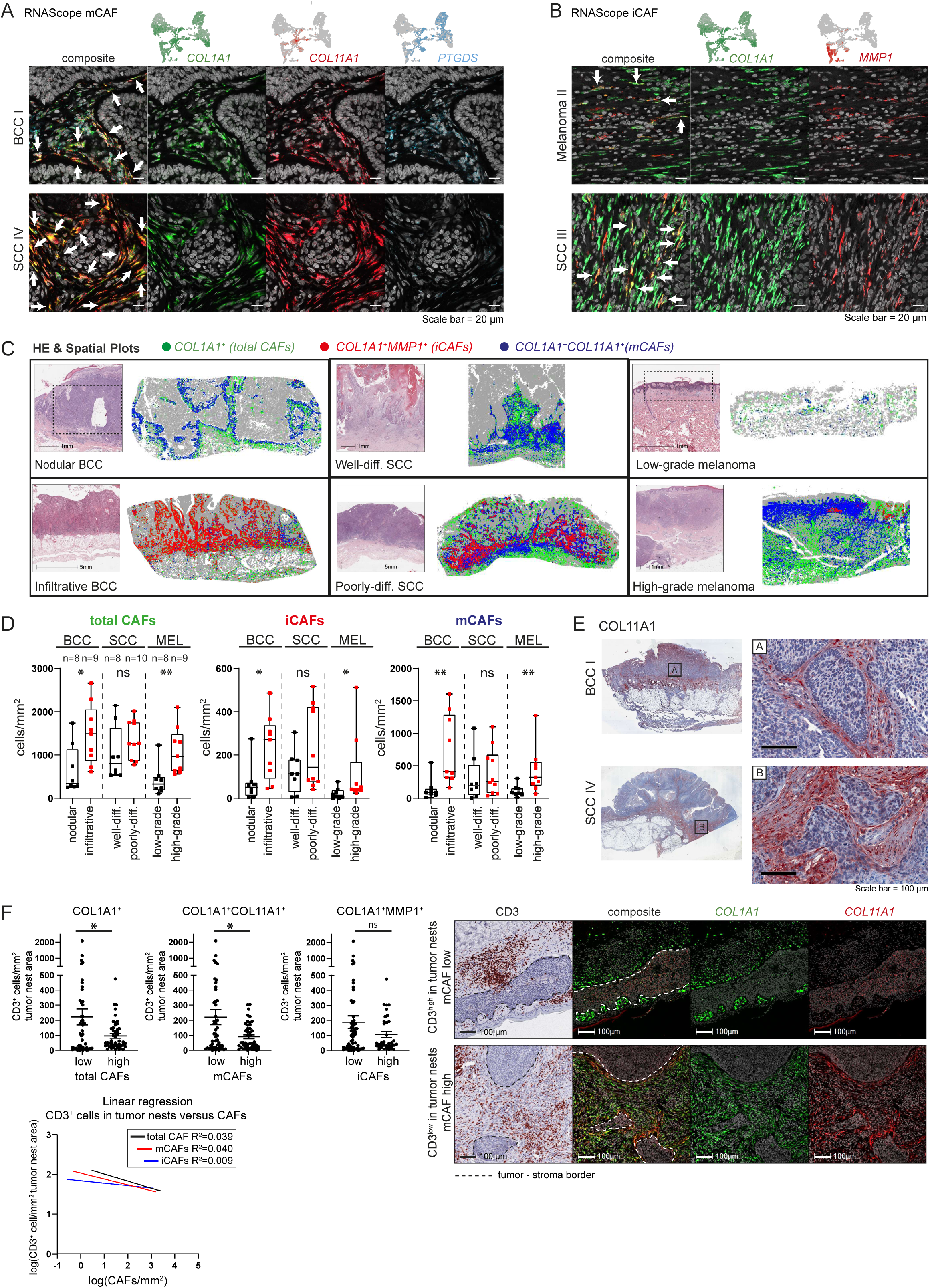
mCAFs and iCAFs are characterized by the expression of ECM and immunomodulatory genes, respectively. (A,B) Representative images from (A) *COL1A1* (green), *COL11A1* (red) and *PTGDS* (blue) and (B) *COL1A1* (green) and *MMP1* (red) RNAScope fluorescence stainings to identifiy mCAFs and iCAFs respectively in FFPE tissue sections from different tumor samples. DAPI nuclear stain is shown in grey. Scale bar represents 20 μm. (B) Spatial plots highlighting the spatial distribution of total CAFs (*COL1A1*), iCAFs (*COL1A1^+^MMP1^+^)* and mCAFs (*COL1A1^+^COL11A1^+^*) and respective H&E stainings on consecutive sections. Dashed-lined boxes show approximate area of spatial plot in H&E staining. (C) Quantification of total CAFs (*COL1A1*^+^), iCAFs (*COL1A1*^+^*MMP1*^+^), and mCAFs (*COL1A1*^+^*COL11A1*^+^*MMP1*^-^) in cells per mm^2^ in 52 samples of nodular (n=8) and infiltrative BCC (n=9), well (n=8) and poorly (n=10) differentiated SCC as well as low-(n=8) and high-grade (n=9) melanoma. Fibroblasts numbers of at least 5 representative ROIs from each tumor were summed-up and normalized to the tissue area to capture the whole tumor tissue. Statistical analysis by Mann Whitney test, p-value ** < 0.01, * < 0.05. (D) Representative images from *COL11A1* immunohistochemistry stainings. Scale bar represents 100 μm. (E) Image analysis of CD3^+^cells/mm^2^ in tumor nests and total CAFs/mm^2^ (high-low cutoff 140 cells/mm^2^), mCAFs/mm^2^ (high-low cutoff 40 cells/mm^2^) or iCAFs/mm^2^ (high-low cutoff 40 cells/mm^2^) in 97 ROIs from nodular and infiltrative BCCs (n=15). Linear regression analysis of log(CD3^+^cells/mm^2^) in tumor nests and log(CAFs/mm^2^). Representative images of CD3 immunohistochemistry and *COL1A1* (green) *COL11A1* (red) RNAScope fluorescence stainings. Statistical analysis by unpaired t-test, *p<0.05.

### mCAFs form a barrier around tumor nests

We investigated the expression of matrix-associated genes (collagens, laminins, lysyloxidases and other ECM genes) and immune response or cancer invasion-associated genes (MMPs, chemokines, interleukins and immunomodulatory genes) by module scores and found a significant enrichment of matrix-associated genes in mCAFs and immuno/invasiveness-associated genes in iCAFs (Figure S8A and S8B). Additionally, we interrogated for possible ligand-receptor interactions: mCAFs exhibit a strong expression of collagens and other ECM genes, whose receptors are found on healthy and neoplastic keratinocytes and melanocytes as well as on immune cells (Figure S9A and S9B). COL11A1 protein staining revealed a dense network of collagen fibers aligning the basement membrane of tumor nests (Figure 5E), indicating that mCAFs may control T cell marginalization as has been shown for COL11A1 expressing CAFs in lung cancer ^72^. Thus, we quantified the number of mCAFs and CD3^+^ cells in the total cancer tissue (including tumor stroma) and the number of CD3^+^ cells within tumor nests of several ROIs per sample of nodular and infiltrative BCC samples (n=15, 97 ROIs). The number of mCAFs/mm² tissue negatively correlated with CD3^+^ cells/mm² within tumor nests (Figure 5F), suggesting that mCAFs may form a physical barrier to inhibit T cell infiltration into tumor nests. Of note, while total CAF and mCAF numbers negatively correlate with CD3 cells/mm² in tumor nests, iCAF numbers did not (Linear regression: total CAFs: R²=0,039; mCAFs: R²=0,040; iCAFs: R²=0,009). Indeed, representative images from overlaid CD3 and mCAF stainings (COL1A1^+^COL11A1^+^) show that infiltration of CD3^+^ cells foremost occurs at areas where staining of mCAFs is low or absent (Figure 5F).

### iCAFs are the major source of cytokines in the TME and are capable of activating T cells

iCAFs strongly express immunomodulatory genes, including *TGFB3* and *LGALS9*, proinflammatory cytokines such as *IL1B* and *IL6* (Figure 6A and S8B). Additionally, iCAFs express high levels of a plethora of chemokines in comparison to healthy or neoplastic keratinocytes and melanocytes (Figure 6A upper heatmap), and thus likely regulate the immune cell composition and influence immune surveillance in the tumor as their receptors are found on many different immune cells. (Figure 6A and 6B). Notably, iCAFs from melanoma samples express high levels of *CXCL1-8* but not *CXCL9-13*, whereas the expression of *CXCL9-13* is high in iCAFs of the SCC III and BCC II sample. Only *CXCL2* is equally high expressed in iCAFs from all tumor samples (Figure 6A). Similarly, *IL1B* is expressed at much higher levels in iCAFs derived from melanoma, while *TGFB3* and *LGALS9* are strongly expressed in iCAFs from BCC and SCC but not melanoma (Figure 6A). We also made a receptor-ligand interrogation with CellChat and confirmed several predicted interaction partners of iCAFs and mCAFs as a signaling source (Figure 6B and S9) ^73^. We further confirmed the CAF-derived expression of cytokines by mRNA stainings *in situ* (Figure 6C, S7B). *CXCL2*, *CXCL8* and *IL24* were selected for the analysis because these three cytokines showed a good coverage across the iCAF cluster, although cell- and sample-specific differences remain (Figure S7B,C). As visualized in the spatial plots of representative samples, cytokine-expressing CAFs are more abundant in the most aggressive tumor variants (infiltrative BCC, poorly differentiated SCCs, and high-grade melanoma) (Figure S7B). While the majority of nodular BCC and low-grade melanomas harbored no or single dispersed cytokine-expressing CAFs, several infiltrative BCC and high-grade melanoma presented with multiple clusters of cytokine-expressing CAFs (Figure 6C), which is in line with the *in situ* quantification of iCAFs (Figure 5C,D) and transcriptomic data (Figure S4A). The difference in cytokine-expressing CAF density and distribution was not as pronounced between well and poorly-differentiated SCCs, although they appeared to be more frequent in late-stage SCC (Figure 6C and S7B). Furthermore, we confirmed that CAFs are a major source for cytokines in the TME in an entirely independent single cell transcriptomics dataset of melanoma (n=5) (Figure S8C). These samples express exceptionally high levels of *CCL2*, *CXCL12* and *CXCL14*. Along these lines, also CAFs from oral SCCs display stronger cytokine expression than their respective tumor cells (Figure S8D).

**Figure 6.**
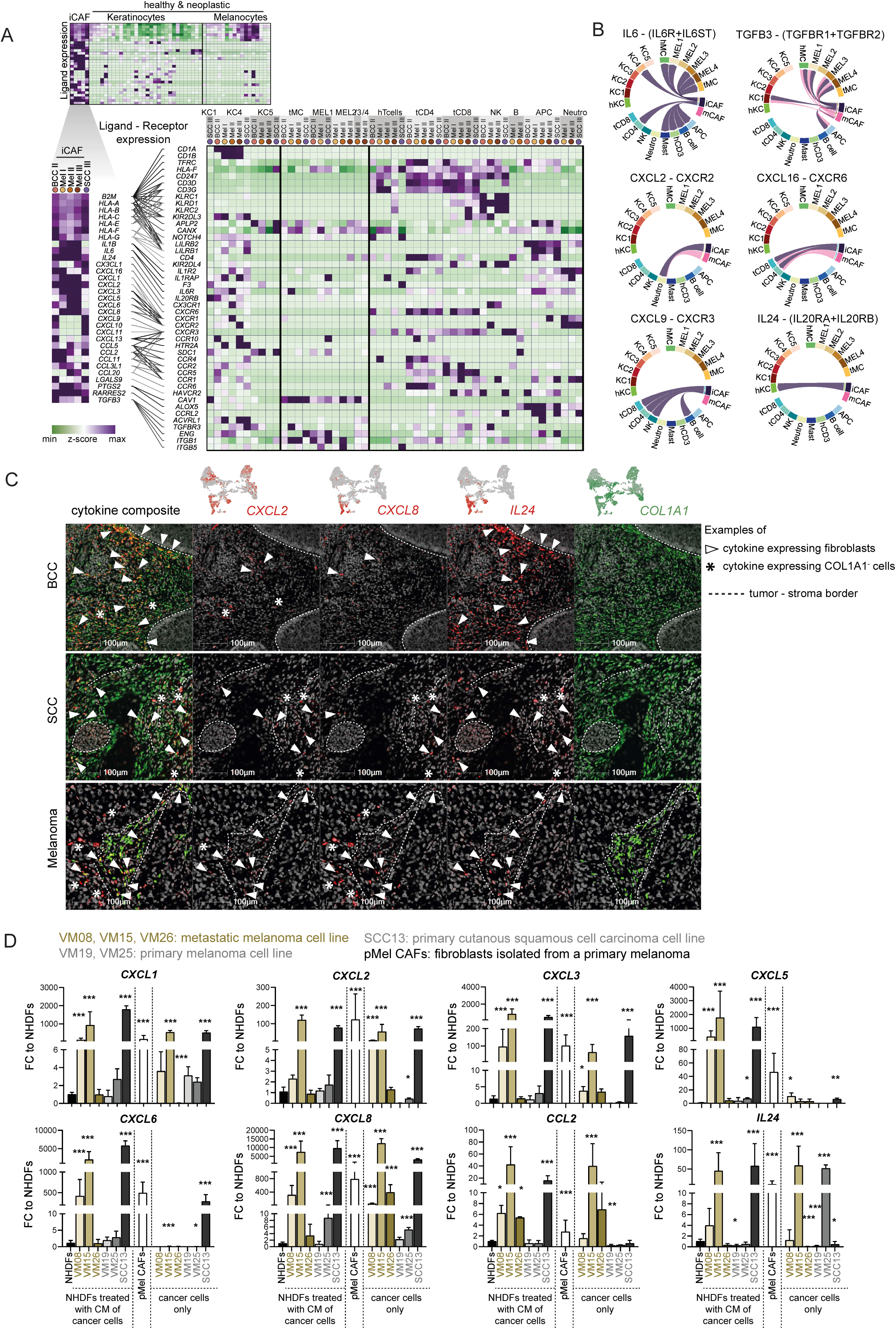
Fibroblasts are an important source of chemokines in the tumor. (A) Expression of immunomodulatory genes in iCAFs compared to healthy and neoplastic keratinocytes and melanocytes and interrogation for respective receptors in healthy and neoplastic keratinocytes and melanocytes as well as immune cells. (B) Circular plots of selected receptor-ligand pairs from CellChat analysis, showing mCAF/iCAF as source cells. (C) Representative images of RNA ISH staining with probes against *CXCL2*, *CXCL8*, *IL24* and *COL1A1* of BCC, SCC and melanoma samples. (D) *In vitro* cytokine expression of NHDF after exposure to conditioned medium from NHDFs, VM08, VM15, VM26, VM19, VM25 and SCC13 cells for 72 hours in comparison to the cytokine expression of the cancer cell lines VM08, VM15, VM19 and SCC13, and to primary melanoma-derived CAFs (pMel CAFs). Statistical analysis by One-way-ANOVA and Tukey’s post hoc test for multiple comparison on log-transformed data. Significant comparisons to NHDFs are shown; *p<0.05, **p<0.01, ***p<0.001.

These results led us to hypothesize that the cancer cells of invasive cancers (but not from non-invasive ones) may directly impact the phenotype of tumor-adjacent fibroblasts. To test this, we isolated primary dermal fibroblasts from healthy skin (NHDF) and treated them with conditioned media collected from melanoma and SCC cell lines (Figure S10A). Intriguingly, the conditioned media of cultured cell lines derived from melanoma metastases (VM08 and VM15 from lymph node metastasis) or from a highly aggressive SCC (SCC13) ^74^ strongly induced the expression of different cytokines and chemokines in healthy skin fibroblasts (Figure 6D). Likewise, these cytokines and chemokines were expressed at high levels by fibroblasts isolated from a primary melanoma without further treatment (pMel CAFs; Figure 6D). On the contrary, VM19 and VM25 cell lines, which were derived from primary melanomas, did not induce cytokine expression (except CXCL8 by VM25). The melanoma cell line VM26 derived from a subcutaneous metastasis ^75^ only induced higher levels of *CCL2* but not the other tested cytokines and chemokines. Although the cancer cells expressed several cytokines themselves (Figure 6D), it was striking that the supernatant of these cancer cell lines induced even more cytokines and chemokines in healthy fibroblasts. Importantly, we confirmed the expression of several cytokines and chemokines by fibroblasts and induced iCAFs on protein level with LEGENDplex assays (Figure S11).

Intriguingly, while the conditioned medium from cancer cells alone induced the expression of iCAF-related genes (cytokines, chemokines, Figure 6D), expression of ECM-related genes was not induced (Figure S10B). Thus, we conclude that the secretome of invasive tumor cell lines can transform normal fibroblasts into iCAF-like, but not mCAF-like cells *in vitro*.

Furthermore, naïve CD4 and CD8 T cells were co-cultured together with NHDFs that were pretreated with conditioned medium from VM15, VM26, VM19, VM25 or control medium, and with pMEL CAFs. In parallel, naïve CD4 and CD8 T cells were co-cultured with the cancer cells directly. We demonstrated that primary fibroblasts isolated from healthy skin are capable of activating T cells (Figure 7A-C and S12A), as shown by increased percentages of proliferating CD4 and CD8 T cells (Figure 7B) and activated CD69+ CD4 and CD8 T cells (Figure 7C, S12A and S12B). This potential to activate T cells was enhanced when fibroblasts were exposed to the secretome of cancer cells. Comparing cancer CM-treated NHDFs to untreated NHDFs, showed a further increase in CD4 T cell proliferation with CM derived from VM15 and VM26, and in CD8 T cell proliferation with CM derived from VM15, VM26 and VM19. Early T cell activation was promoted by VM15- and VM19-derived CM for CD4 T cells, and by VM15-derived CM for CD8 T cells (Figure 7C). Late activation of CD4+ T cells measured as percentage of CD45RO+/CD62L-T cells at 96h was significantly enhanced by CM derived from VM15, VM26 and VM19 (Figure S12C and S12D). Importantly, also CAFs directly isolated from a primary melanoma without further treatment (pMel CAFs) were potent in activating CD4 and CD8 T cells (Figure 7B,C).

**Figure 7.**
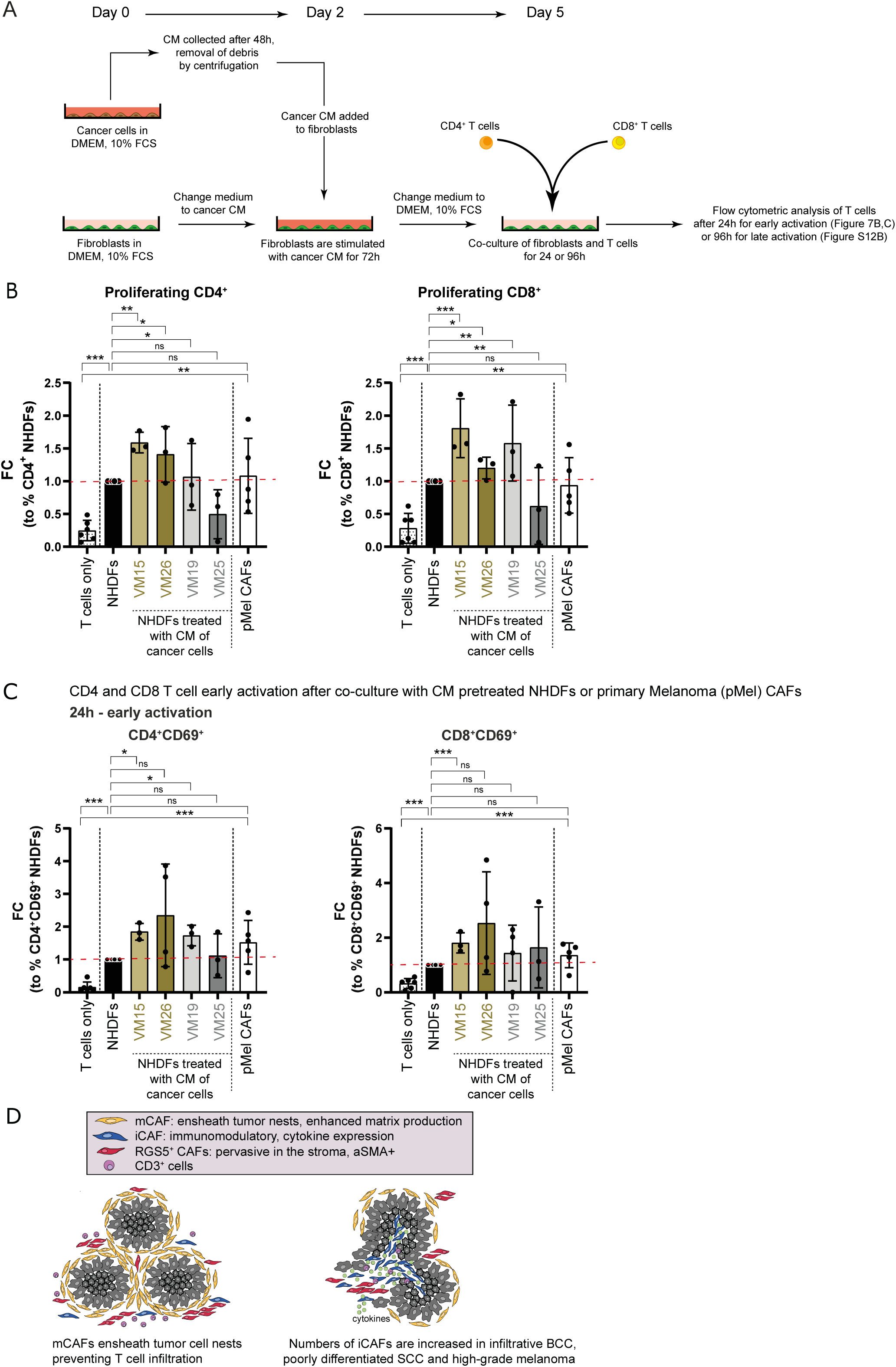
Fibroblasts activate CD4^+^ and CD8^+^ T cells. (A) Experimental setup of T cell assays shown in B and C. (B) Proliferation assessed by flow cytometry of CD4 or CD8 T cells upon co-culture with NHDFs pre-treated with conditioned medium from cancer cells, primary melanoma-derived CAFs (pMel CAFs) or cancer cells. (C) Upregulation of the early activation marker CD69 on CD4 or CD8 T cells after 24h of co-culture with NHDFs pre-treated with conditioned medium from cancer cells, pMel CAFs or cancer cells. Data represented as fold change of percentages of cells positive for the indicated markers normalized to NHDFs. (B,C) Statistical analysis in comparison to NHDFs or to T cells only by unpaired Student’s t test; *p<0.05, **p<0.01, ***p<0.001. (D) Schematic summary of spatial distribution of distinct CAF subsets in human skin cancer.

Taken together, *in situ* stainings of the marker genes identified in our scRNAseq screen showed that mCAFs and iCAFs are distinct CAF populations that follow different distribution patterns *in situ* (Figure 7D). mCAFs are present in all tumors but seem to play an important role at the tumor-stroma border as they form dense networks surrounding tumor nests of benign tumors, i.e. nodular BCC and well differentiated SCC. In contrast, the number of iCAFs increases significantly in aggressive tumors (especially infiltrative BCC and late-stage melanoma) and, thus, high abundance of iCAFs correlates with malignant progression (Figure 5D and 7D). Importantly, receptor-ligand analysis revealed that iCAFs, which are associated with late-stage tumors, express many immunomodulatory factors that bind to receptors expressed primarily on neutrophils, T cells and NK cells. Notably, the heat map in Figure 6A shows that apart from immune cells, fibroblasts synthesize the majority of cytokines and chemokines but not tumor cells, indicating that not the cancer cells but the stromal cells (fibroblasts) are key players in immune cell recruitment and activation. Indeed, we confirmed that fibroblasts treated with the secretome of skin cancer cells are capable of activating T cells. Furthermore, since mCAFs ensheath tumor nests and synthesize large amounts of ECM proteins, it is likely that they are involved in T cell exclusion (Figure 5E,F).

## Discussion

Fibroblasts are important contributors to the TME. They can exert pro- as well as anti-tumorigenic functions by stimulating tumor cell survival and proliferation, modifying ECM stiffness, supporting metastasis, influencing therapy response, regulating immune cell recruitment via chemokine secretion and inflammatory responses ^36,76^. Inter- and intratumoral CAF heterogeneity have been appreciated ever since scRNA-seq methods have become available. However, their different functions in the TME remain largely inexplicit. The scRNA-seq based works that have been published for skin cancers, four melanoma studies ^21,23–25^, one SCC study ^22^ and one study containing BCCs and SCCs ^26^, either only contained a very small number of fibroblasts or fibroblast heterogeneity was not the focus of the analysis. Considering the large knowledge about fibroblast diversity in healthy human dermis ^10–12,39^, a screen on the CAF heterogeneity in human skin tumors was required to fill the missing gap, which we achieved with the present work, especially since our dataset comprises the three major skin cancer types, and because we validated our findings not only in other published datasets but also by comprehensive spatial analysis of the different CAF subsets *in situ*.

We have previously shown that healthy human skin comprises two functionally distinct fibroblast subsets (papillary and reticular fibroblasts) which can be distinguished by the expression of *CD90* (*THY1*) ^39^. The present scRNA-seq screen confirmed that *CD90*, which is still frequently used as the sole fibroblast marker to isolate or visualize skin fibroblasts, is only expressed by the reticular subpopulation (Figure S4C). Although the majority of CAFs express *CD90* (Figure S4C), our data do not allow to conclude whether CAFs develop only from skin-resident reticular fibroblasts or whether they acquire expression of *CD90* upon activation.

Of note, the RNA expression level of *FAP* in the scRNA-seq data does not reflect the FAP protein expression on the fibroblasts entirely, as we used FAP and CD90 surface expression to enrich for fibroblasts by FACS (Figure S1B and S4C).

In the skin cancer samples, we identified three distinct fibroblast populations (excluding the small population of ucCAFs). Immunomodulatory CAFs (iCAFs) show a characteristic expression of proinflammatory cytokines (*IL1B*, *IL6*), chemokines (CXCR2 ligands) and immunomodulating molecules (*IDO1*) and thus seem to be analogue to the previously described iCAFs in other cancer types (Figures 3B, 5E and 5F) ^35^. Matrix CAFs (mCAFs), which we identified as a separate CAF population, are not to be confused with myofibroblasts described in other publications^35^. mCAFs exhibit increased matrix production but without the expression of *ACTA2* and myosin light chain proteins (Figures 3B, S8A and S8B).

Most interestingly, our dataset suggested that iCAFs and mCAFs can likely be attributed to skin cancer types with higher (iCAFs) and lower (mCAFs) metastatic potential. The only exception were fibroblasts from BCC II, which were found in the iCAF cluster even though BCC II is histologically not different to the other two BCC samples (Figure S3). Quantification of the distinct CAF subsets was particularly challenging because of the varying nature of skin cancer. Skin tumors display heterogeneous morphology both within a single tumor (i.e. superficial tumor areas versus invasive front) and among distinct cancer subtypes. Thus, the biological difference may not be well captured in numbers. Large-field spatial visualization of the CAF subsets in tumor tissue samples showed an obvious difference in the CAF patterns from lower to higher malignancy, along with a higher overall CAF density in the more aggressive variants of the respective skin cancer subtypes (Figure 5C,D). *In situ* localization revealed that mCAFs are present in all tumors but are detected in high density at the tumor-stroma border especially in nodular BCC and well-differentiated SCC. That mCAFs also increase in number in late-stage tumors (Figure 5, S6A and S7) is interesting in connection with the results of our multiplex mRNA staining *in situ*. Future spatial transcriptomic analysis could reveal if the mCAFs located at the tumor-stroma border have a distinct expression profile compared to the mCAFs located within the stromal strands and without direct contact to tumor cells. This may indicate, for example, dual functions of mCAFs or a potential conversion of tumor ensheathing-CAFs towards a “more aggressive” iCAF-like phenotype both of which are an exciting future route to explore.

iCAFs are more abundant in malignant tumors, especially in infiltrative BCC and in aggressive melanomas (Figure 5B,C). Although there is a trend that poorly differentiated SCCs harbor higher numbers of iCAFs compared to well-differentiated SCCs, the difference is not so clear between these two cancer subtypes. Analysis of larger patient cohort with additional stratification into further subtypes may be necessary to provide a clearer picture.

The *RGS5^+^* cluster contains cells from all samples, including healthy controls. According to differential gene expression, the *RGS5^+^* cluster would commonly be termed as myofibroblasts/myoCAFs due to the characteristic expression of *ACTA2*, *MYH11* and *COL1A1*. However, the *RGS5^+^*cluster also showed a marker profile that can be attributed to pericytes (*RGS5*, *PDGFRB*, *KCNJ8*, *TAGLN, MCAM*) (Figure 3B and 4A). We demonstrate by *in situ* immunohistochemistry that *RGS5^+^* cells are not only found in a perivascular localization but are also distributed throughout the stroma in tumor regions. In contrast, in unaffected skin, *RGS5^+^* cells are restricted to a perivascular localization (Figure 4A and 4B). Thus, an exclusive definition as myofibroblasts/myoCAFs or pericytes for this cluster seems to be inappropriate. The impossibility to discern myofibroblasts/myoCAFs from pericytes on RNA level has been a general issue, as we found very similar expression patterns of the myofibroblasts/pericytes in a HNSCC dataset, once defined as myofibroblasts ^19^, and once defined as pericytes ^71^ (Figure 4C). It has been suggested that pericytes are able to leave the vessel wall and contribute to the tumor stroma in a process called pericyte-fibroblast transition (PFT) ^77^. To our knowledge, PFT has not been shown in skin cancer. A recent pan-cancer scRNA-seq study suggested that a small CAF subset arose from endothelial cells. However, this was concluded from transcriptional data but not confirmed *in situ* ^78^. Whether *RGS5*^+^ CAFs originate from pericytes or skin-resident fibroblasts cannot be concluded from this study. It is of higher importance that *RGS5^+^ACTA2^+^*CAFs are present in all tumor samples, which might represent a general phenotype that is similar to activated fibroblasts expressing *ACTA2* in non-cancerous conditions, such as wound healing ^79,80^. Notably, Grout et al. recently described an *MYH11*^+^*aSMA*^+^(*ACTA2*^+^) and *COL4A1* expressing CAF subset in non-small cell lung cancer that might be involved in T cell exclusion. The *RGS5^+^* CAFs in our study (which we appointed a T cell-exclusion role from tumor nests) also expressed *MYH11*, *ACTA2* and *COL4A1*^72^.

Reanalysis of published datasets from HNSSC ^18^ and cutaneous SCC ^21^ confirms the presence of RGS5^+^ CAFs in both SCC types (Figure S13A and B), and revealed CAF subsets with expression patterns similar to mCAFs and iCAFs in cutaneous SCC (Figure S13B, CAF1 and CAF2). In HNSCC, the CAF1 subset displayed expression of both mCAF and iCAF genes (Figure S13A). However, in the cutaneous SCC dataset, CAFs from one patient are overrepresented (>50% of total fibroblasts; 92% of CAF1 and 70% of CAF2 subsets), which impedes to deconstruct if CAF1 or CAF2 are more or less abundant in moderate and well differentiated SCCs. We also confirmed the presence of all three CAF subsets in an independent melanoma dataset (Figure S7C). Moreover, reanalysis of single cell transcriptomic data of 5 infiltrative BCCs^27^ confirmed the presence of mCAFs and a small cluster of iCAFs (Figure S13C). Furthermore, the samples included a fibroblast cluster expressing signature genes of reticular fibroblasts (Figure S13C). Of note, also in our dataset few cells from BCC and SCC contributed to the rFIB cluster, most of them however from unaffected skin adjacent to the tumors (Figure S4A). However RGS5^+^ cells were not included in their CAF population but might be part of the pericyte population, which clustered separately in their first level clustering ^27^. Garnier et al. identified two clusters of *RGS5*+ and *TAGLN*+ pericytes in healthy skin and BCCs ^53^. They detected a selective expansion of *RGS5*+ pericytes and a reduction in *TAGLN*+ pericytes in BCC compared to healthy skin, and described that the colocalization with vessel-like structures is lost in BCC, indicating that these cells are similar to the *RGS5*+ *TAGLN*+ CAFs described in our study. Furthermore, they detected four fibroblasts subsets which were designated as APOD+, *SFRP2*+, *PTGDS*+, and *POSTN*+. Intriguingly, both *POSTN*+ and *PTGDS*+ CAFs were detected around the tumor islands, suggesting that these CAFs correspond to the mCAFs in our study as we used *COL11A1* and *PTGDS* to localize them in the tissue (Figure 5A).

CAFs share common features with fibroblastic reticular cells (FRCs) within lymph nodes, which generate ECM conduits to guide the traffic of immune cells and the transit of potential antigens ^81^. It is established that CAFs participate in T-cell exclusion from tumor nests ^82^. Several studies have reported reduced T cell infiltration in CAF-rich tumors compared to their CAF-low counterparts ^83^. mCAFs are detected at the tumor-stroma interface and ensheath tumor nests, especially in nodular BCC and well differentiated SCC. As they synthesize a range of ECM proteins including COL11A1, which we detected as dense fibers surrounding tumor nests, we propose that mCAFs play a crucial role in T cell marginalization. Indeed, we detected a negative correlation between T cell numbers present in tumor nests and the number of mCAFs surrounding the tumor nests (Figure 5F). A similar function was described for a subset of FAP^+^ αSMA^+^ lung CAFs expressing COL11A1/COL12A1 or COL4A1 in human lung cancer ^72^. Thus, targeting mCAFs may improve the efficacy of immunotherapy in patients bearing T cell-excluded tumors. Indeed, a whole tumor cell vaccine genetically modified to express FAP significantly reduced cancer growth in a murine model of lung cancer and melanoma by directly inhibiting CAFs and simultaneously enhancing T cell infiltration ^84^. Whether the ECM barrier formed by mCAFs modulates marginalization of other immune cells or inhibits or promotes tumor cell invasion, remains to be explored.

The importance for immunomodulatory chemokines in cancer progression is undisputable. The expression of CXCR2 ligands, *CXCL1-3* and *CXCL5-8* by melanoma cells has been shown to control the immune cell composition of the TME, contribute to the ability to escape tumor immune surveillance, induce angiogenesis or define the preferred sites of melanoma metastases ^85–88^. In the present study, receptor-ligand analysis revealed that fibroblasts are the major source for cytokines and chemokines (Figure 6A,B) and not the cancer cells themselves, thus highlighting the importance of fibroblasts in immune cell recruitment and cancer immune surveillance.

Intriguingly, while *CXCL2* was expressed by fibroblasts from all skin cancer types, melanoma-derived CAFs expressed high levels of *CXCL1-3, 5, 6* and *8* and *IL1B* as well as *IL6*, whereas the expression of *CXCL9-11* and *13* was high in non-melanoma CAFs (Figure 5E). *LGALS9*, which has been shown to interact with *CD40* on T cells thereby attenuating their expansion and effector function, was strongly expressed in CAFs from BCC and SCC but not melanoma. Furthermore, HLA genes were highly expressed in CAFs but not normal fibroblasts (Figure S8B), suggesting a role for CAFs as antigen-presenting cells. CAF-mediated cross-presentation of neo-antigens may directly suppress T cell function ^89^. These findings indicate that although iCAFs are present in melanoma and non-melanoma skin cancers, the expression of chemokines and possibly other immunomodulating genes is tumor type-dependent. This is also reflected by the differentially regulated expression of TFs in iCAFs derived from melanoma and cSCCs (Figure 3C). We substantiated the presence of differential crosstalk among tumor types and stages by testing the effect of conditioned media from various cancer cell lines on healthy skin-derived fibroblasts. Indeed, conditioned medium from metastasis-derived melanoma cell lines induced an iCAF-like phenotype and cytokine/chemokine expression while conditioned medium of a primary melanoma cell line did not change the cytokine/chemokine expression (Figure 6D). Surprisingly, the secretome of the subcutaneous metastasis-derived cell line VM26 ^75^ did not induce the expression of the majority of the tested cytokines and chemokines except for *CCL2*, which may be linked to different mutations. However, VM26-derived conditioned medium was still capable of activating T cells, which is not surprising as the cytokines not induced by VM26 shown in Figure 6D are CXCR1/2 ligands that are known to recruit innate immune cells but not T cells. Interestingly, while the secretome of melanoma and SCC cell lines was capable of inducing an iCAF phenotype, induction of a mCAF phenotype could not be achieved *in vitro.* Thus, further investigations are necessary to define which signals or culture conditions prime fibroblasts towards mCAF differentiation *in vitro* and *in situ*. Likewise it remains elusive whether soluble CAF-derived factors or direct cell contact are essential for T cell activation.

In addition to the fibroblast heterogeneity in skin tumors, our data highlight the tremendous effect of the TME on all cells within a tumor. For example, melanocytes from BCC and SCC samples (tMC), which are part of the non-neoplastic cells in these tumor types, cluster separately from melanocytes that were derived from healthy skin samples (hMC) (Figure 2B and S2C). This indicates that the altered gene expression profile is likely induced by the TME. Further, our CNV analysis clearly shows that samples from the same tumor subtype (Melanoma I – III: Acral lentiginous melanoma, ALM) and body location can greatly differ at molecular level, which explains donor-specific clustering.

In summary, our work provides a cellular and molecular atlas of the three most frequent skin cancer types comprising neoplastic epithelial, mesenchymal, and immune cells. We further reveal and characterize three distinct CAF subsets and show that their abundance and associated signaling molecules and structural proteins critically impact the TME. Therefore, determining the predominant CAF subset within tumor samples may improve future diagnostic strategies and thereby open new avenues for better personalized therapies. Moreover, pharmacologically targeting CAFs to reduce ECM density could enhance T cell trafficking into tumor nests and thus the efficacy of checkpoint inhibition therapy as well as the penetration of agents that directly target cancer cells.

## Methods

### Human healthy skin and tumor samples

Fresh 4 mm punch biopsies from central tumor and unaffected skin adjacent to tumors as well as 10x10 cm healthy skin samples from abdominal plastic surgeries were subjected to cell isolation procedure directly after surgery.

Healthy skin samples III-IV were cut into thin strips after removal of the fat layer. Epidermis was separated from dermis by Dispase 2 (1:100, Roche #04942078001, 20mg/mL), digested in PBS at 37°C for one hour before being peeled off. The epidermal sheet was minced and then subjected to enzymatic digestion in Trypsin-EDTA (GIBCO #25300-054) for 20 min at 37°C in a shaking water bath (Epidermal sheet protocol for enrichment of keratinocytes). Healthy skin dermis and tumor samples were cut into tiny pieces and digested with Collagenase 1 (1:100, GIBCO #17100-017, 50mg/mL), Collagenase 2 (1:100, GIBCO #17101-015, 50mg/mL), Collagenase 4 (1:100, Sigma-Aldrich #C5138, 50mg/mL), Hyaluronidase (1:100, Sigma-Aldrich #H3884, 10mg/mL) and DNAseI (1:250, Sigma-Aldrich #DN25, 5 mg/mL) in DMEM/10%FCS for one hour in a 37°C water bath (Protocol for enrichment of fibroblasts, keratinocytes and immune cells). After enzymatic digestion, the cell suspension was filtered and washed in PBS/10% FCS twice before subjecting it to FACS staining.

After Fc blocking (1:500, CD16/CD32 BD #553142, RRID:AB_394656), cell suspensions were stained for 30 min at 4°C in the dark with CD45-BV605 (1:50, BioLegend #304042, RRID:AB_2562106), ITGA6-PeCy7 (1:100, BioLegend #313622, RRID:AB_2561705), CDH1-PeCy7 (1:200, BioLegend #147310, RRID:AB_2564188), FAP-APC (1:20, R&D Systems #FAB3715A, RRID:AB_2884010), CD90-AF700 (1:30, BioLegend #328120, RRID:AB_2203302) and CD31-FITC (1:30, BD Biosciences #563807), CD106-Pacific Blue (1:100, BD Biosciences #744309, RRID:AB_2742138), CD235ab-Pacific Blue (1:1000, BioLegend #306611, RRID:AB_2248153) and DAPI. ITGA6^+^/CDH1^+^ keratinocytes, FAP^+^/CD90^+^ fibroblasts, CD45^+^ immune cells and FAP^-^CD90^-^ double negative cells were single cell sorted directly into Smart-seq2 lysis buffer in 384-well plates. After sorting, plates were stored at -80°C until they were sent for sequencing to the Eukaryotic Single Cell Genomics Facility (ESCG) at SciLifeLab at the Karolinska Institutet, Sweden.

### Ethical approval

Written informed patient consent was obtained before tissue collection in accordance with the Declaration of Helsinki. Our study was approved by the Institutional Review Board under the ethical permits EK#1695/2021, EK#1783/2020 and EK#1555/2016.

### Immunohistochemistry

Immunohistochemistry was performed on 4μm human FFPE sections according to standard protocols. Antigen retrieval was conducted in citrate buffer, pH 6.0 and 3%BSA/PBST was used for blocking. Primary antibodies against CD90 (THY1) (1:200, rabbit monoclonal [EPR3133], Abcam #ab133350, RRID:AB_11155503), FAP (1:200, rabbit monoclonal [D3V8A], Cell signaling #13801, RRID:AB_2798316), TAGLN (1:12.000, rabbit polyclonal, Thermo Scientific #PA5-27463, RRID:AB_2544939), DES (1:2000, rabbit monoclonal [Y266], Abcam #ab32362, RRID:AB_731901), CD31 (1:500, rabbit, Neomarkers, #RB10333-P1)), CD3 (1:200, rabbit, Abcam #ab16669, RRID:AB_443425) and COL11A1 (1:200, rabbit polyclonal, Abcam #ab64883, RRID:AB_1140613) were diluted in 1%BSA/PBST and incubated over night. A biotinylated goat anti-rabbit antibody (1:200, Vector BA-1000) was used as second step and incubated for 30 min at room temperature. Novocastra Streptavidin-HRP (Leica Biosystems Newcastle #RE7104) and Dako AEC+ High sensitivity substrate (Dako #K3469) were used for signal enhancement and development. For counter staining, hematoxylin was used.

### RNAScope

RNAScope was conducted by using Multiplex Fluorescent Reagent Kit v2 from Advanced Cell Diagnostics, ACD Bio-Techne (#323135), according to the manufacturer’s protocol, with probes for *COL1A1* (#401891-C2), *COL11A1* (#400741-C3), *PTGDS* (#431471-C1), *MMP1* (#412641-C1), *RGS5* (#533421-C3) and *PDGFRA* (#604481-C1). For fluorescence staining, the TSA dyes Fluorescein, Cy3 and Cy5 (Akoyabio) and DAPI as nuclear stain were utilized. The Multiplex Fluorescent Reagent Kit v2 and the RNAscope® 4-Plex Ancillary kit were combined for 4-plex stainings against CXCL2 (#425251), CXCL8 (310381-C2), IL24 (404301-C3) and Col1A1 (401891-C4), or against MMP1 (#412641-C1), Col11A1 (#400741-C2), RGS5 (#533421-C3) and COL1A1 (#401891-C4). Opal™ fluorophores (Opal 520 or Opal780, Opal 570, Opal 620 and Opal 690) were utilized in the 4-plex staining’s. Images were captured by Vectra Polaris™ and image analysis was conducted with HALO® image analysis platform.

### Quantification of total CAFs, iCAFs and mCAFs in tumor sections

To analyze and quantify the various CAF populations in tumor sections, we utilized the HALO® image analysis platform. The cell types included total CAFs (COL1A1^+^), iCAFs (COL1A1^+^MMP1^+^), and mCAFs (COL1A1^+^ COL11A1^+^ MMP1^-^), which were identified by specific marker combinations. Tumor sections from nodular (n=8) and infiltrative BCC (n=9), well (n=8) and poorly (n=10) differentiated SCC, as well as low (n=8) and high (n=9) grade melanoma were analyzed. A minimum of five representative regions of interest (ROIs) per sample were selected, excluding scarred or ulcerated areas to ensure accurate quantification. The HALO v3.6.4134 software with the HighPlex FL v4.2.14 plugin was employed for image analysis, with signal intensity thresholds set for each channel to differentiate positive from negative cells. Representative spatial plots were generated from analyzed sections in HALO®.

### T cell exclusion from tumor nests

BCC samples (n=15) were stained for immunohistochemistry (IHC) with antibodies against CD3, followed by staining of consecutive sections using the RNAscope 4-Plex Multiplex Fluorescent Reagent Kit to *COL1A1*, *MMP1*, *Col11A1* and *RGS5*. HALO® image analysis platform was used to perform image analysis and quantification. Regions of interest (ROI) were selected, where CD3^+^ cells were present in the stroma surrounding the tumor nests. A machine learning classifier, which was then trained to differentiate between tumor and stromal tissue, was applied to the ROIs. We quantified the number of CD3^+^ cells in the total tissue area or only in the tumor nests within the ROI. The sections were then co-registered with the RNAscope staining, and the numbers of CAFs (*COL1A1*^+^ cells) and matrix CAFs (*COL1A1*^+^*COL11A1*^+^*MMP1*^-^*RGS5*^-^) were quantified. The cell counts were then normalized to cells/mm².

### Fibroblast activation by conditioned medium of cancer cells: transcriptomic and proteomic analysis

Normal healthy dermal fibroblasts (NHDF) and fibroblasts from a primary melanoma were isolated as described above and cultured in DMEM containing 10% FBS and 50 µg/ml Gentamycin in a humidified incubator at 37°C and 5% CO_2_.

### Generation of conditioned medium (CM) from cancer cell lines

Melanoma cell lines (VM08, VM15, VM19, VM25, VM26)^75^ were cultured in RPMI1640 (GIBCO #11875093) containing 10% FBS (GIBCO #26140079), 2 mM L-glutamine (GIBCO #25030081) and 50 U/ml streptomycin/penicillin (GIBCO #15070063). SCC13^74^ cell line (RRID:CVCL_4029) was cultured in DMEM Glutamax containing 10 % FBS, 2 mM L-glutamine (GIBCO #25030081), 50 U/ml streptomycin/penicillin (GIBCO #15070063), 5µg/ml insulin and 10 µg/ml transferrin. When cells reached 70-80% confluency, they were washed with PBS, and DMEM/10% FBS was added. Conditioned medium (CM) was collected 48 hours later, centrifuged with 300 g for 10 minutes and stored at -20°C.

### Fibroblast activation assay

NHDF and cancer cells were seeded into 6 well plates for 24 hours. Then, medium of the NHDF was exchanged with CM derived from cancer cells or from NHDF as a control. Cells were harvested 72 hours later. RNA was isolated with the Qiagen RNeasy Mini Kit (Qiagen #74106). RevertAid H Minus First Strand cDNA Synthesis Kit (Thermo Scientific #K1631) was used to prepare cDNA after a DNaseI digestion step (Thermo Scientific #EN0521). Taqman 2xUniversal PCR Master Mix (Applied Biosystems #4324018) and Taqman probes for GAPDH (Hs99999905), CXCL1 (Hs00236937_m1), CXCL2 (Hs00601975_m1), CXCL3 (Hs00171061_m1), CXCL5 (Hs01099660_g1), CXCL6 (Hs00605742_g1), CXCL8 (Hs00174103_m1), CCL2 (Hs00234140_m1) and IL24 (Hs01114274_m1), Lumican (Hs00929860_m1), Col11A1 (Hs01097664_m1), Col4A1 (Hs00266237_m1), LOXL2 (Hs00158757_m1), Fibromodulin (Hs05632658_s1) and Col12A1 (Hs00189184_m1) were used in the qPCR.

#### LEGENDplex Assay

For proteomic analysis cancer CM or control medium was removed from fibroblasts after 72h incubation. Cells were washed in PBS and fresh DMEM/10% FCS was added for 48h. Supernatants were collected, centrifuged to remove cell debris and subjected to protein analysis using LEGENDplex kits (Human Essential Immune Response #740930 and Human Proinflammatory Chemokine Panel #740985, BioLegend). Data analysis was done in GraphPad Prism version 8.0.0 for Windows, GraphPad Software, San Diego, California USA, www.graphpad.com.

### T cell activation assay

#### Naïve T cell isolation

Human peripheral blood obtained from healthy individuals (with informed consent), was collected in heparinized tubes and immediately processed by mixing 1:1 with PBS, then layered over Ficoll-Paque™ PLUS (Cytiva). After density gradient centrifugation at 500 x g, 20°C, for 20 minutes, the PBMC layer was transferred, washed with PBS, and CD4+ and CD8+ T cells were isolated using the human CD4+ and CD8+ T cell isolation kits (Miltenyi Biotec), according to the manufacturer’s protocol. The isolated CD4+ and CD8+ T cells were stained with a mix of antibodies for 20 minutes at 4°C in the dark: CD4-FITC (1:400, [RPA-T4], BioLegend #300501, RRID:AB_314070), CD8-PE-Cy7 (1:100, [HIT8a] BD Biosciences Cat# 555635, RRID:AB_395997), CD25-APC (1:100, [BC96], BioLegend Cat# 302609 (also 302610), RRID:AB_314279), CD14-APC-Cy7 (1:100, [M5E2], BioLegend Cat# 301820 (also 301819), RRID:AB_493695), CD45RO-PacificBlue (1:100, [UCHL1], BioLegend Cat# 304215 (also 304216), RRID:AB_493658), CD127-PE (1:100, [A019D5], BioLegend Cat# 351340 (also 351303, 351304), RRID:AB_2564136), CD16-PerCP-Cy5.5 (1:500, [3G8], (BioLegend Cat# 302028 (also 302027), RRID:AB_893262)). Cells were washed with PBS and resuspended in RPMI1640 without phenol red (Gibco) containing a 1:1000 dilution of 7-AAD viability dye (BioLegend). Naïve CD4 or CD8 T cells were sorted with the BD FACSAria™ II Cell Sorter based on CD4+CD127+CD14-CD16-CD45RO-CD25- or CD8+CD127+CD14-CD16-CD45RO-CD25-. The collected T cells were stained with a proliferation dye (1:500, eBioscience, V450) for 8 minutes at 37°C, followed by washing with RPMI1640 media (Gibco) containing 10% FBS, 1% penicillin/streptomycin (Gibco), and 2 mM L-glutamine (Gibco).

#### Experimental setup and FACS analysis

2000 fibroblasts/CAFs or melanoma cells were seeded into 96-well plates using DMEM or RPMI1640 media (Gibco), containing 10% FBS, 1% penicillin/streptomycin (Gibco), and 2 mM L-glutamine (Gibco), respectively. After 24 hours, the medium of NHDFs were replaced with conditioned medium derived from either NHDFs or cancer cells. Following 72h incubation at 37°C, cells were washed with PBS, and 50 µL of RPMI containing 10% FBS, 1% penicillin/streptomycin (Gibco), 2 mM L-glutamine and 12.5 µL/mL Immunocult CD3/CD28 T cell activator (Stemcell) was added. Then 50 µL of RPMI containing 40,000 T cells at a ratio of 1 (CD8) to 1.5 (CD4) were added. The cells were harvested by scraping after 24h or 96h and subjected to FACS analysis for the assessment of T cell proliferation (24h) and expression of CD69 (24h), CD45RO (96h) and CD62L (96h) on CD4 or CD8 T cells (negative of fibroblast markers CD90 and FAP). Cells were stained in PBS/10%FCS and incubated for 20 minutes at 4°C in the dark: CD4-FITC (1:400, [RPA-T4], BioLegend #300501, RRID:AB_314070), CD8-PE-Cy7 (1:100, [HIT8a] BD Biosciences Cat# 555635, RRID:AB_395997), CD69-PE (1:100, [FN50], BioLegend Cat# 985202, RRID:AB_2924641), CD90-AF647 (1:100, [5E10], BioLegend Cat# 328115 (also 328116), RRID:AB_893439), FAP-APC (1:100, [427819], R&D Systems Cat# FAB3715A-025), CD45RO-PacificBlue (1:100, [UCHL1], BioLegend Cat# 304244 (also 304205, 304206), RRID:AB_2564160), CD62L-PerCP-Cy5.5 (1:100, [DREG56], Elabscience Cat# E-AB-F1051A). Following incubation, cells were washed and resuspended in PBS with Fixable Viability Dye eFluor™ 780 APC-Cy7 (eBioscience) for subsequent analysis using CytoFlex LX Flow Cytometer (Beckman Coulter).

### Single cell RNA sequencing

scRNA-seq was conducted by the Eukaryotic Single Cell Genomics Facility (ESCG) at SciLifeLab, Sweden according to the Smart-seq2 protocol ^90^. Demultiplexed reads were aligned to the human genome (hg19 assembly) and the ERCC spike-in reference using STAR v2.4.2a in two-pass alignment mode ^91^. Uniquely aligned reads were transformed into reads per million kilobase (RPKM) using rpkmforgenes(). RPKM values were summed up when several isoforms of a gene were detected.

### Single cell RNA sequencing data processing

Low quality cells were removed by using thresholds for RPKM-values and number of genes expressed per cell. The lower threshold was referred to empty-well controls, the upper threshold was set based on unusually high RPKM-values of clearly visible outliers. We considered good quality when a minimum of 400 genes and RPKM-values between 150.000-8.000.000 per cell was reached. Finally, 4824 cells were retained after quality control. Subsequent data analysis was carried out by using R3.6.2 and the Seurat package v3 (Stuart*, Butler*, et al., Cell 2019). RPKM values for each gene per cell were normalized and natural-log transformed (NormalizeData: normalization.method=”Log-Normalize”). The 2000 most variable genes were identified (FindVariableGenes: selection.method = ‘vst’), the data scaled and principal component analysis (PCA) was performed. The first 20 principal components 1:20, resolution 0.2 and Seurat default parameters were used for UMAP generation of first-level clustering. Subsequently, clusters for second-level clustering were selected based on commonly known signature gene expression: Healthy keratinocytes, SCC and BCC (*KRT5*, *KRT14*), melanocytes and melanoma cells (*MLANA*), immune cells (*CD45*) as well as fibroblasts and vSMCs (*COL1A1*, *RERGL*).

Differentially expressed genes were identified by the Seurat function FindAllMarkers. For generation of UMAPs, violin and bar plots, ggplot2 v3.3.2 were used.

### Copy number variations for estimation of malignancy

*InferCNV* of the Trinity CTAT Project (https://github.com/broadinstitute/inferCNV) was used to calculate CNVs for healthy and malignant keratinocytes and separately for melanocytes and melanoma cells in comparison to stromal cells. For CreateInfercnvObject, healthy stromal cells (Healthy donor cells from Fibroblast&vSMC second level clustering, which includes fibroblasts, vSMCs and pericytes) were used as reference for CNV estimation. *InferCNV* operations were performed by infercnv::run using min_cells_per_gene = 3, cutoff = 1, cluster_by_groups = T, denoise = T, HMM = T, analysis_mode = subclusters, hclust_method = ward.D2, tumor_subcluster_partition_method = random_trees. Estimation of malignancy was performed as previously described ^25^, using a pearson correlation cutoff of 0.45 or 0.40 and a sum of squares (SoS) cutoff of 0.017 or 0.026 for healthy and malignant keratinocytes or melanocytes and melanoma cells, resepctively. In the CNV estimation plots for melanocytes and melanoma cells (Figure S3B) we highlighted melanocytes derived from cluster hMC (melanocytes derived from healthy skin) and tMC (melanocytes derived from SCC, BCC and melanoma adjacent from unaffected skin samplesadjacent to melanoma) where most of the cells are nicely found in the lower quadrants as expected (CNV- and undefined). CNV estimation based on RNA expression only detects genomic aberrations that affects larger chromosomal sections. Thus, we also analyzed the expression of certain genes in keratinocytes from BCC and SCC samples in comparison to healthy keratinocytes. For determining neoplastic keratinocytes in BCC samples we used PTCH1 and PTCH2^55,56^ (Figure S2A,B).

### Trajectory analysis

We used Monocle2^65^ (v2.28, R4.0.0) and Monocle3^66^ (v0.2.1, R3.6.2) to perform trajectory analysis. For both methods, we extracted RPKM data, phenotype data, and feature data from the Seurat object (second-level clustering of fibroblasts without vSMC) from which we created a newCellDataSet(lowerDetectionLimit = 0.1, expressionFamily = tobit()) or a new_cell_data_set() object using default parameters.

For Monocle2, we converted our RPKM data into mRNA counts using relative2abs() and generated the NewCellDataSet(lowerDetectionLimit = 0.5, expressionFamily = negbinomial.size()) object again. As quality filtering and clustering were already performed in Seurat, we directly constructed single cell trajectories using all significantly (adjusted p-value < 0.01) regulated DEGs (FindMarkers()) as input parameters for ordering cells. For calculating pseudotime, we used healthy skin cells from controls as our starting point. Cells were plotted using plot_cell_trajectory() colored by “clusters”, “category” and “pseudotime”.

For Monocle 3, we manually added clusters, UMAP and PCA parameters to the new_cell_data_set() object and calculated the trajectory graph with learn_graph(object, use_partition = F). For calculating pseudotime we used healthy skin clusters (pFIB and rFIB) as root_cells and used plot_cells(color_cells_by = “pseudotime”) to present the data.

### Heatmaps

*ComplexHeatmap* v2.2.0 function was used to represent gene expression of single cells or mean gene expression per cluster in heatmaps as z-scores.

### Receptor-Ligand Analysis

For receptor-ligand pairing the previously published method developed by Simon Joost was used, but with additional adjustments for run-time and parallel computing ^93^. Receptor-ligand interactions were analyzed between fibroblast clusters, immune cell clusters, neoplastic and healthy keratinocyte and melanocyte clusters.

A signature gene list, containing potential ligands and receptors of each cluster, was generated by the Seurat function FindMarkers() at the level of second-level clustering.

Potential ligand-receptor interactions were identified by querying the combined receptor-ligand database from Ramilowski et al., 2015 and Cabello-Aguilar et al., 2020.

For each cluster pair, the number of identified receptor-ligand pairs was compared to the number of pairs obtained from an equally sized randomly sampled pool of receptors and ligands. This was repeated 10.000 times to test for significantly enriched interactions (p ≤ 0.05 for Benjamini-Hochberg-corrected p-values). An additional prerequisite for a valid receptor-ligand pairing was the presence of at least 2.5% cells of the same donor in both of the potentially interacting clusters (eg. iCAFs interacting with tCD4 requires at least 2.5% cells from the same donor in each of the clusters).

Used packages: *python* 3.7.6, *pandas* 1.0.1, *numpy* 1.18.1, *matplotlib* 3.2.2. Receptor – Ligand heatmaps were generated with *seaborn* 0.11.0 using mean z-scores per donor per cluster.

Additionally, we verified receptor-ligand interactions with CellChat ^73^. The communication probability was calculated according to default parameters. We present selected receptor-ligand pairs as circular plots using the function netVisual individual(source.use = c(“mCAF”, “iCAF”), layout = chord).

### Module Score

Module Scores were calculated by AddModuleScore() function from Seurat, and genes sets were represented as violin plots. Individual genes of a gene set are shown in heatmaps. Genes that showed absolutely no expression in any cluster were excluded from heatmaps and module score calculation (chemokines: CCL4, CCL14, CXCL7; cytokines: IL9, IL31; MMPs: MMP26). Statistical analysis was done by non-parametric Wilcox rank-sum test using ggplot2 function stat_compare_means().

### Melanoma scRNAseq Validation Dataset

For the melanoma validation dataset, pre-treatment samples (n=5) were collected from stage IV melanoma patients as part of a trial investigating anti-CD20 treatment in a therapeutic setting (10.1038/s41467-017-00452-4). After biopsy of a lesion, single cell suspensions were immediately frozen. Thawed suspensions were subjected to scRNA-seq using the Chromium Single Cell Controller and Single Cell 5’ Library & Gel Bead Kit v1.1 (10X Genomics, Pleasanton, CA) according to the manufacturer’s protocol. Sequencing was performed using the Illumina NovaSeq platform and the 150bp paired-end configuration.

Preprocessing of the scRNA-seq data was performed using Cell Ranger version 6.1.2 (10x Genomics). Expression data was processed using R (version 4.2.1) and Seurat (version 4.0.5). Cells with less than 1,000 genes or more than 10% of relative mitochondrial gene counts were removed. The data was processed following Seurat’s scTransform workflow. Sample-specific batch effects were corrected using Harmony. Clusters were identified using Seurat’s “FindNeighbors” and “FindClusters” functions, using the first 32 dimensions of the Harmony-corrected embedding and a resolution of 1.5. Cell types were subsequently identified based on canonical markers. Heatmaps and module scores were generated as described above.

### Publicly available dataset

We reanalyzed the publicly available HNSCC data set (GSE103322) using R3.6.2 and the Seurat package v3 ^19^. As described by the authors, we regressed for the variable *processedbyMaximaenzyme*. The cell annotation, which was provided in the meta data file, was used to select the cells for clustering of the fibroblasts. Based on markers that were described by the authors, the clusters for CAFs, myofibroblasts and intermediate fibroblasts were assigned. A heatmap (ComplexHeatmap v2.2.0) was generated presenting gene expression as means of z-scores per cluster using the same genes as shown in the heatmap in Figure S2C of the original publication, but extended it by commonly accepted pericyte and vSMC markers.

Additionally, we identified our marker genes for mCAFs, iCAFs, RGS5^+^ cells and healthy fibroblasts in the fibroblast population of the HNSCC data set (GSE103322)^19^, cutaneous human SCC (GSE144240)^22^ and human invasive BCC (GSE181907)^27^ and represented it in heatmaps showing gene expression as means of z-scores.

For the human cutaneous SCC (GSE144240) dataset, fibroblast cell annotation was provided by the authors.

For the human invasive BCC data set, cell annotations for the fibroblast subclusters (FC1-FC4), as described in the original paper ^27^ were provided by the authors upon request. As some marker genes were expressed in very few cells, we used a cutoff of at least 2% of cells expressing the gene in order to include it into the heatmap.

For reanalysis of this datasets we used R3.6.2, Seurat package v3 and *ComplexHeatmap* v2.2.0.

### Data availability

Raw data are available at the European Genome-Phenome Archive (EGAS50000000365) and expression matrices are accessible at GEO (GSE254918).

## Acknowledgements

We cordially thank Hao Yuan, Alexander Enz and Matthias Wielscher for assisting with bioinformatic data processing, Robert Loewe and Philipp Weber for their help in sample collection, Robin Ristl from the Institute of Medical Statistics for his support in statistical data analysis, as well as Erwin Wagner, Reinhard Kirnbauer and Alessandra Handisurya for their support and critical feedback. Special thanks to Nina Zila, Monika Weiss and Karin Neumüller for their technical assistance and lab organization. We thank Andreas Spittler, Günther Hofbauer as well as Marion Gröger, Sabine Rauscher and Christoph Friedl from the Flow Cytometry and Imaging core facilities of the Medical University Vienna for their continuous support. Finally, we thank the Eukaryotic Single Cell Genomics Facility (ESCG) at SciLifeLab, Sweden for sequencing our cells and the national UPPMAX SENS computational infrastructure for data transfer. Computation of *inferCNV* and *CellChat* were performed using the Vienna Scientific Cluster 3 (VSC-3).

