## Supplemental Information for "Cancer associated fibroblast subtypes modulate the tumor-immune microenvironment and are associated with skin cancer malignancy"

#### SUPPLEMENTAL TABLES AND FIGURES:

**Table S1.** Published skin cancer scRNA-seq datasets including fibroblasts/CAFs

**Table S2.** Details of donor samples

**Figure S1.** Gating scheme and quality control of sequencing data. Related to Figure 1

**Figure S2.** Estimation of malignant cells and additional second-level clustering information. Related to Figure 2

**Figure S3.** Estimation of malignant cells by Pearson correlation and sum of squares (SoS) of CNVs according to Tirosh et al., 2016. Related to Figure 2.

**Figure S4.** Characterization of the fibroblasts and vSMC cluster. Related to Figure 3.

**Figure S5.** Endothelial and vCAF markers in the *RGS5*+ cluster. Related to Figure 4.

**Figure S6.** RNAScope staining for mCAFs and iCAFs. Related to Figure 5.

**Figure S7.** Spatial analysis of CAF subtypes *in situ*. Related to Figure 4, 5 and 6.

**Figure S8.** Matrix-associated, MMP and immunomodulatory gene panels characterize mCAFs and iCAFs. Related to Figures 5 and 6.

**Figure S9.** Receptor-Ligand analysis of mCAFs. Related to Figure 5.

**Figure S10.** Expression of ECM genes is not upregulated in NHDFs upon treatment with tumor CM. Related to Figure 6.

**Figure S11.** Protein expression in supernatants of fibroblasts upon stimulation with cancer cell-derived CM. Related to Figure 6.

**Figure S12.** Fibroblast-mediated late activation of CD4 and CD8 T cells. Related to Figure 7.

**Figure S13.** Reanalysis of published datasets reveals CAF subsets with gene signatures similar to mCAFs, iCAFs and $RGS5^{+}$ cells in human oral and cutaneous SCC as well as invasive BCC.

#### SUPPLEMENTAL TABLES:

**Supplemental Table 1.** Published skin cancer scRNA-seq datasets including fibroblasts

| Manuscript | cancer type | specimen | total cells | CAFs | additional information |
|---|---|---|---|---|---|
| Tirosh et al., Science 2016 | melanoma | 19 | 4,645 | 61 | |
| Jerby-Arnon et al., Cell 2018 | melanoma | 33 | 7,186 | 106 | |
| Ji et al., Cell 2020 | SCC | 10 | 48,000 | 882 | >50% of CAFs from 1 donor |
| Guerrero-Juarez et al., Science Advances 2022 | BCC | 4 | 36,392 | 5,775 | CAFs only from 2 donors |
| Yerly et al., Nature Communications 2022 | BCC | 5 | 28,819 | 809 | |

**Supplemental Table 2.** Details of donor samples

| Donors | Sample type | Body Part | Sex | Age at sampling (yrs) | Histology |
|---|---|---|---|---|---|
| BCC I | BCC | capillitium | male | 47 | Nodular basal cell carcinoma |
| BCC II | BCC | nose | male | 80 | Nodular basal cell carcinoma |
| BCC III | BCC | shoulder | male | 77 | Nodular basal cell carcinoma |
| Mel I | Melanoma | heel | male | 78 | Acral lentiginous melanoma, T4b, thickness 12 mm |
| Mel II | Melanoma | toe | male | 86 | Acral lentiginous melanoma, T4b, thickness 8 mm |
| Mel III | Melanoma | toe | female | 87 | Acral lentiginous melanoma, T4b, thickness 5 mm |
| SCC I | SCC | capillitium | male | 56 | Reoccurrence of poorly differentiated SCC in scar, thickness 3 mm |
| SCC II | SCC | lower leg | male | 78 | Well differentiated SCC, thickness 2 mm |
| SCC III | SCC | capillitium | male | 93 | Poorly differentiated SCC, thickness 4 mm |
| SCC IV | SCC | lower leg | male | 92 | SCC arisen from Bowens disease thickness 8 mm |
| Healthy I | Healthy skin | upper arm | female | 63 | |
| Healthy II | Healthy skin | abdomen | female | 43 | |
| Healthy III | Healthy skin | abdomen | female | 59 | |
| Healthy IV | Healthy skin | upper arm | female | 44 | |
| Healthy V | Healthy skin | abdomen | male | 48 | |

**SUPPLEMENTAL FIGURE LEGENDS:**

**S1. Gating scheme and quality control of sequencing data. Related to Figure 1.**
(A) Gating strategy for keratinocytes from healthy skin epidermal sheets (Healthy III - V).
(B) Gating strategy for fibroblasts, immune cells, double negatives and keratinocytes of BCC, SCC, melanoma and healthy skin dermis. Immunohistochemistry for FAP and CD90 in healthy skin dermis and tumor samples.
(C) Quality control of scRNA-seq data: Number of genes (left) and RPKM values (right) per sample after quality filtering.
(D) Number of cells per donor sample after quality filtering.

**S2. Estimation of malignant cells and additional second-level clustering information. Related to Figure 2.**
(A) *PTCH1* and *PTCH2* expression levels (RPKM) in healthy and malignant keratinocytes shown per cluster and per donor. Threshold for overexpressing cells was adjusted to exclude healthy keratinocytes (*red dashed line: 100 RPKM*).
(B) *GLI1, GLI2* and *MYCN* expression levels in healthy and malignant keratinocytes shown per cluster and per donor.
(C) UMAPs showing the distribution of donor samples per cluster, bar plot showing the total cell number and distribution of sample category per cluster.

**S3. Estimation of malignant cells by Pearson correlation and sum of squares (SoS) of CNVs according to Tirosh et al., 2016. Related to Figure 2.**
(A) Malignancy estimation for healthy and malignant keratinocytes. Cells, which overexpress *PTCH1* and *PTCH2*, but do not show CNVs are highlighted in green.
(B) Malignancy estimation for melanocytes and melanoma cells. Cells from the healthy melanocyte cluster (hMC) and the tumor melanocyte cluster (tMC) are highlighted in green. Upper right quadrant shows CNV+ cells, upper left quadrant shows intermediate cells, lower left quadrant shows CNV- cells and lower right quadrant shows unclassifiable cells.

**S4. Characterization of the fibroblasts and vSMC cluster. Related to Figure 3.**
(A) UMAP showing the distribution of donors, sample type, sample category and sex on fibroblasts and vSMC second-level clustering.
(B) Bar plots showing the distribution of fibroblasts and vSMC clusters among donor samples (top) or number of fibroblasts and vSMC per donor samples (bottom).
(C) Commonly accepted papillary and reticular markers in the healthy fibroblast clusters pFib and rFib.
(D) Expression of previously described CAF markers on second-level clustering of fibroblasts and vSMCs.

**S5. Endothelial and vCAF markers in the *RGS5*$^{+}$ cluster. Related to Figure 4.**
(A) Previously described vCAF marker genes in a murine model for breast cancer are expressed in the *RGS5*$^{+}$ cluster.
(B) Absence of expression of endothelial cell marker genes in the *RGS5*$^{+}$ cluster.
(C) *COL1A1* (green), *RGS5* (red) and *PDGFRA* (blue) RNAScope fluorescence stainings in BCC II. Scale bar represents 20 $\mu$m.

**S6. RNAScope staining for mCAFs and iCAFs. Related to Figure 5.**
(A) Representative images of RNAScope fluorescence stainings from BCC and SCC samples: *COL1A1* (green), *COL11A1* (red), DAPI (grey). Dashed line showing the tumor-stroma border.
(B) Representative images of RNAScope fluorescence stainings from BCC, SCC and Melanoma: *COL1A1* (green), *COL11A1* (blue), *MMP1* (red) and DAPI (grey).

**S7. Spatial analysis of CAF subtypes *in situ.* Related to Figures 4, 5 and 6.**
(A) Representative images of RNAScope fluorescence stainings for iCAFs, mCAFs and RGS5$^+$cells: *MMP1, COL11A1* and *RGS5* shown as single stainings (*red*) or in combination with *COL1A1 (green), MMP1 (blue)*, *COL11A1 (red) or RGS5 (green)* in composite images.
(B) Spatial plots highlighting the spatial distribution of total CAFs (*COL1A1*) and cytokine expressing CAFs (co-expression of *COL1A1* and *CXCL2, CXCL8* or *IL24).*
(C) *CXCL2, CXCL8* or *IL24* expression in fibroblasts of BCC, SCC or melanoma samples that contribute to the iCAF cluster.

**S8. Matrix-associated, MMP and immunomodulatory gene panels characterize mCAFs and iCAFs. Related to Figures 5 and 6.**
(A) Collagens, Laminins, Lysyl oxidases and other ECM proteins as well as (B) matrix metalloproteinases, chemokines, cytokines and immunomodulatory molecules represented in heatmaps and corresponding Violin plots showing module scores.
(C,D) Expression of cytokines, chemokines and immunoregulatory molecules in fibroblasts or tumor cells of melanoma (C; n=5) or HNSCC (D; dataset of Puram et al., 2018) represented in heatmaps and corresponding Violin plots showing module scores. UMAP in (C) shows identified CAF subsets in melanoma. Statistical analysis by Wilcoxon test, reference group mCAFs (A), iCAFs (B) or tumor cells (C,D), p-value ** < 0.01, * < 0.05.

**S9. Receptor-Ligand analysis of mCAFs. Related to Figure 5.**
(A) Expression of ECM genes in mCAFs compared to healthy and neoplastic keratinocytes and melanocytes (top heatmap) and interrogation for corresponding receptors in healthy and neoplastic keratinocytes and melanocytes as well as immune cells.
(B) Circular plots of selected receptor-ligand pairs from CellChat analysis, showing mCAF/iCAF as source cells for ligand expression.

**S10. Expression of ECM genes is not upregulated in NHDFs upon treatment with cancer cell-derived CM. Related to Figure 6.**
(A) Experimental setup for assessing transcriptional changes in NHDFs after stimulation with cancer-cell derived CM.
(B) *In vitro* ECM expression of NHDFs after exposure to conditioned medium from NHDFs, VM15, VM26, VM19 and VM25 for 72 hours in comparison to CAFs isolated from a primary melanoma (pMel CAFs) and to the cancer cell lines VM15, VM26, VM19 and VM25. Statistical analysis by One-way-ANOVA and Tukey`s post hoc test; Significant comparisons to NHDFs are shown; *p<0.05, **p<0.01, ***p<0.001.

**S11. Protein expression in supernatants of fibroblasts upon stimulation with cancer cell-derived CM. Related to Figure 6.**
(A) Experimental setup for generation of supernatants of NHDFs stimulated with cancer cell-derived CM.
(B) Protein levels (pg/mL) of several cytokines and chemokines assessed by LegendPlex. Statistical analysis by One-way-ANOVA and Tukey`s post hoc test; Significant comparisons to NHDFs are shown; *p<0.05, **p<0.01, ***p<0.001.

**S12. Fibroblast-mediated late activation of CD4 and CD8 T cells. Related to Figure 7.**
(A) Gating strategy for detection of early and late activation in CD4 and CD8 T cells including FMO controls for CD69, CD62L and CD45RO.
(B) Percentage of CD4$^+$CD69$^+$ and CD8$^+$CD69$^+$ T cells in co-culture with control NHDFs.

(C) Late activation ($CD45RO^{+}CD62L^{-}$) of CD4 or CD8 T cells after 96h of co-culture with CM pre-treated NHDFs or cancer cells. Statistical analysis in comparison to NHDFs or to T cells only by unpaired Student's t test; *p<0.05, **p<0.01, ***p<0.001.
(D) Percentage of $CD4^{+}CD45RO^{+}CD62L^{-}$ and $CD8^{+}CD45RO^{+}CD62L^{-}$ in co-culture with control NHDFs.

**S13. Reanalysis of published datasets reveals CAF subsets with gene signatures similar to mCAFs, iCAFs and $RGS5^{+}$ cells in human oral and cutaneous SCC** as well as invasive BCC.
(A) UMAPs, heatmaps of top DEG from mCAFs, iCAFs and $RGS5^{+}$ cells and module scores of chemokines, immunomodulators, collagens and other ECM molecules of CAFs in human oral SCC (Puram et al.,2017).
(B,C) UMAPs and heatmaps of top DEG from mCAFs, iCAFs, $RGS5^{+}$ cells and healthy fibroblasts from human cutaneous SCC (Ji et al., 2022) (B) or BCC (Yerly et al., 2022) (C).

### Figure S1

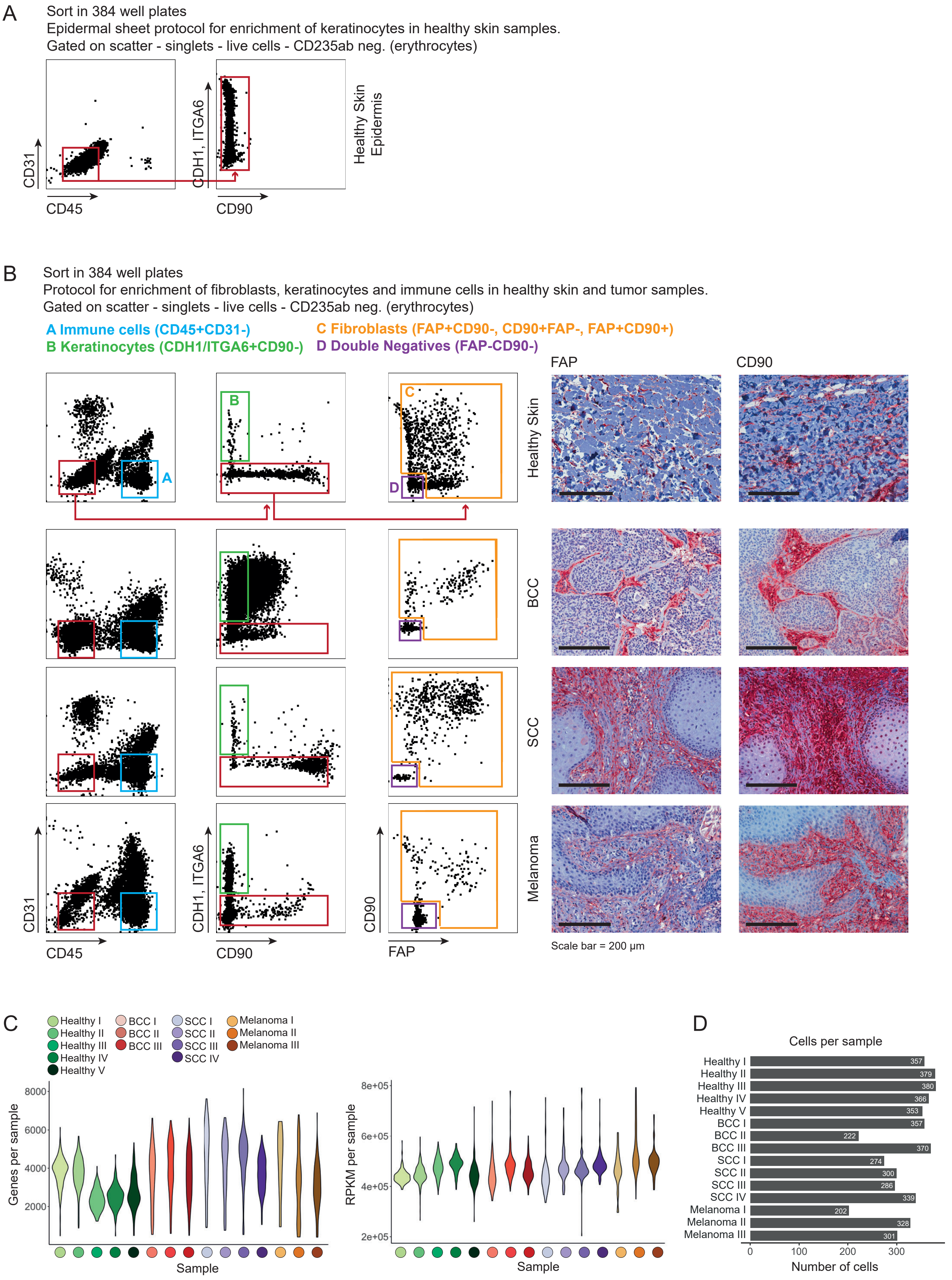
A
Sort in 384 well plates
Epidermal sheet protocol for enrichment of keratinocytes in healthy skin samples.
Gated on scatter - singlets - live cells - CD235ab neg. (erythrocytes)
CD31
CD45
CDH1, ITGA6
CD90
Healthy Skin Epidermis
B
Sort in 384 well plates
Protocol for enrichment of fibroblasts, keratinocytes and immune cells in healthy skin and tumor samples.
Gated on scatter - singlets - live cells - CD235ab neg. (erythrocytes)
A Immune cells (CD45+CD31-)
B Keratinocytes (CDH1/ITGA6+CD90-)
C Fibroblasts (FAP+CD90-, CD90+FAP-, FAP+CD90+)
D Double Negatives (FAP-CD90-)
FAP
CD90
Healthy Skin
BCC
SCC
Melanoma
CD31
CD45
CDH1, ITGA6
CD90
CD90
FAP
Scale bar = 200 µm
C
Healthy I
Healthy II
Healthy III
Healthy IV
Healthy V
BCC I
BCC II
BCC III
SCC I
SCC II
SCC III
SCC IV
Melanoma I
Melanoma II
Melanoma III
Genes per sample
8000
6000
4000
2000
Sample
RPKM per sample
8e+05
6e+05
4e+05
2e+05
Sample
D
Cells per sample
Healthy I 357
Healthy II 379
Healthy III 380
Healthy IV 366
Healthy V 353
BCC I 357
BCC II 222
BCC III 370
SCC I 274
SCC II 300
SCC III 286
SCC IV 339
Melanoma I 202
Melanoma II 328
Melanoma III 301
0
100
200
300
Number of cells

Figure S2

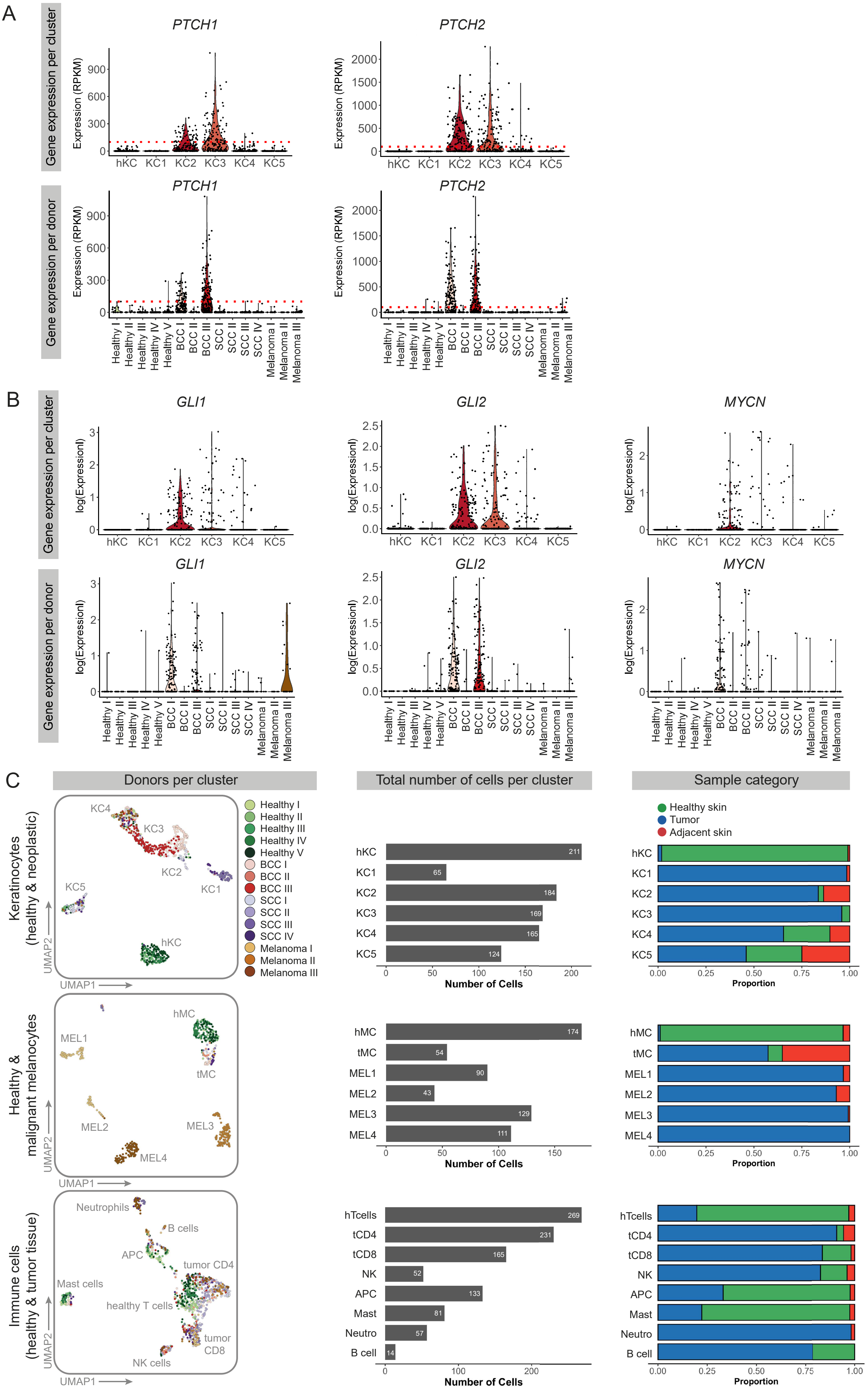

A
Gene expression per cluster
PTCH1
PTCH2
Expression (RPKM)
hKC
KC1
KC2
KC3
KC4
KC5
Gene expression per donor
Healthy I
Healthy II
Healthy III
Healthy IV
Healthy V
BCC I
BCC II
BCC III
SCC I
SCC II
SCC III
SCC IV
Melanoma I
Melanoma II
Melanoma III
B
GLI1
GLI2
MYCN
log(Expression)
C
Donors per cluster
Total number of cells per cluster
Sample category
Keratinocytes (healthy & neoplastic)
KC4
KC3
KC2
KC1
KC5
hKC
UMAP1
UMAP2
Healthy I
Healthy II
Healthy III
Healthy IV
Healthy V
BCC I
BCC II
BCC III
SCC I
SCC II
SCC III
SCC IV
Melanoma I
Melanoma II
Melanoma III
hKC 211
KC1 65
KC2 184
KC3 169
KC4 165
KC5 124
Number of Cells
Healthy skin
Tumor
Adjacent skin
Proportion
Healthy & malignant melanocytes
hMC
MEL1
tMC
MEL2
MEL3
MEL4
hMC 174
tMC 54
MEL1 90
MEL2 43
MEL3 129
MEL4 111
Immune cells (healthy & tumor tissue)
Neutrophils
B cells
APC
tumor CD4
Mast cells
healthy T cells
tumor CD8
NK cells
hTcells 269
tCD4 231
tCD8 165
NK 52
APC 133
Mast 81
Neutro 57
B cell 14

### Figure S3

A

Estimation of copy number variations (CNVs)
Healthy and malignant keratinocytes

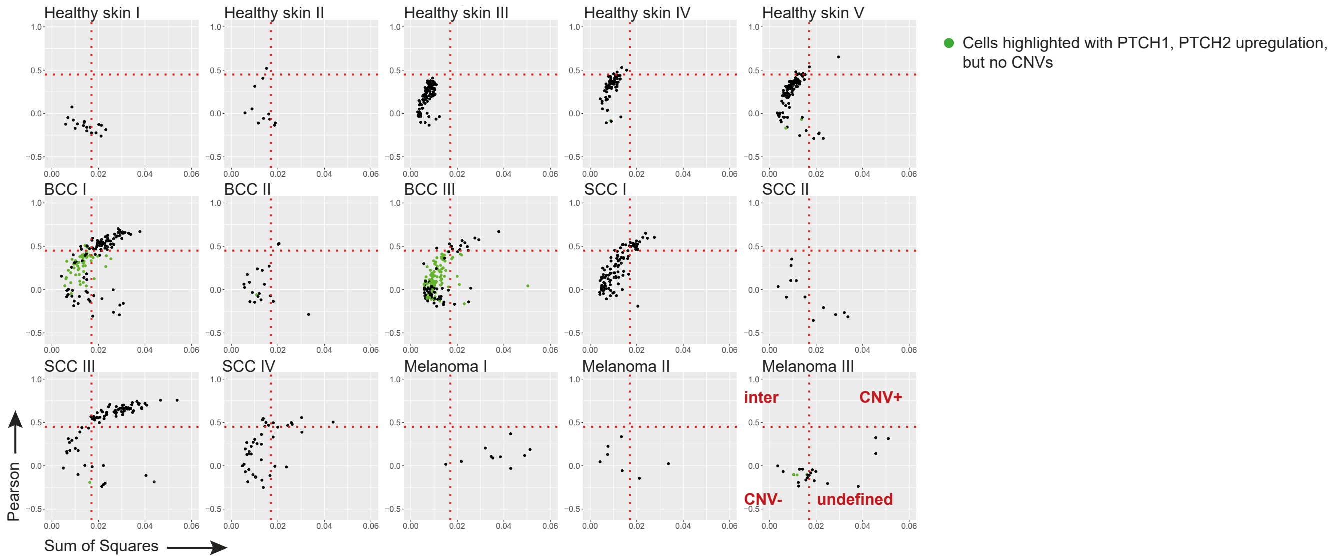


B

Estimation of copy number variations (CNVs)
Melanocytes and melanoma cells

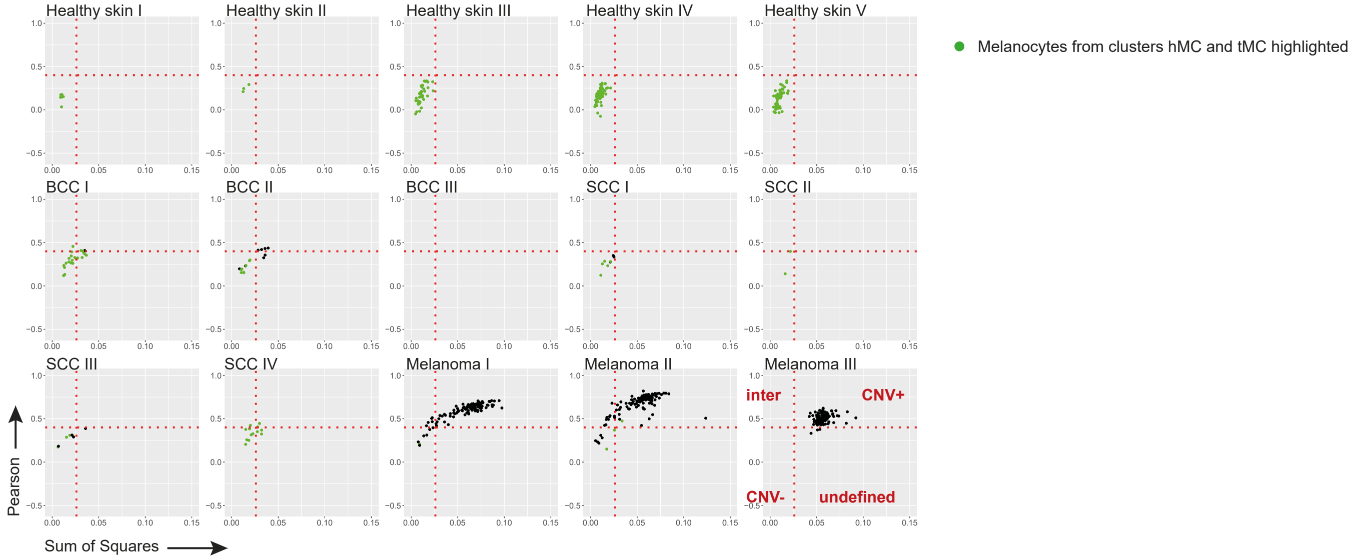

Figure S4

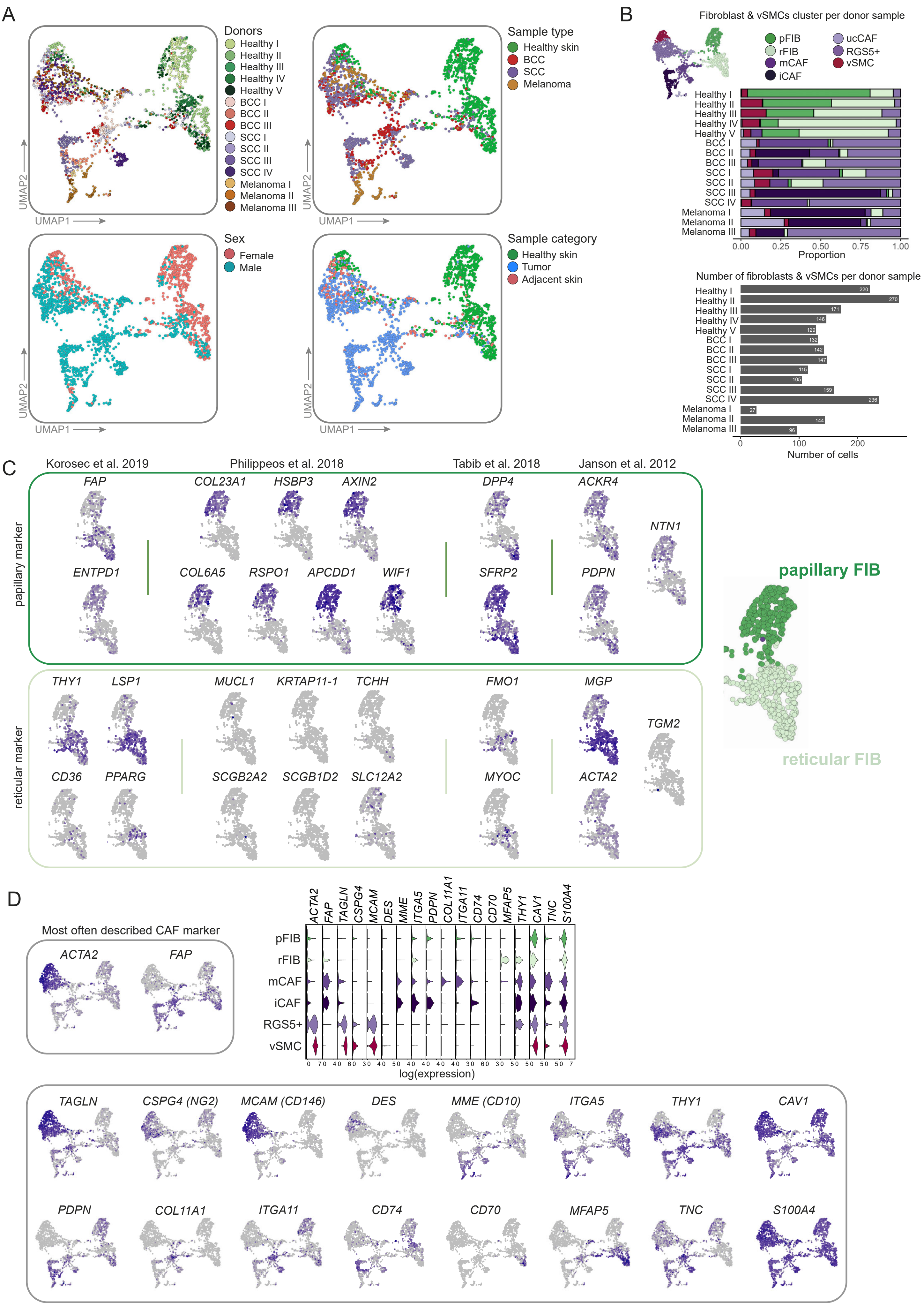


Reviewed in Nurmik et al. 2020, Han et al. 2020, Gascard et al. 2016, Xing et al. 2011

Figure S5

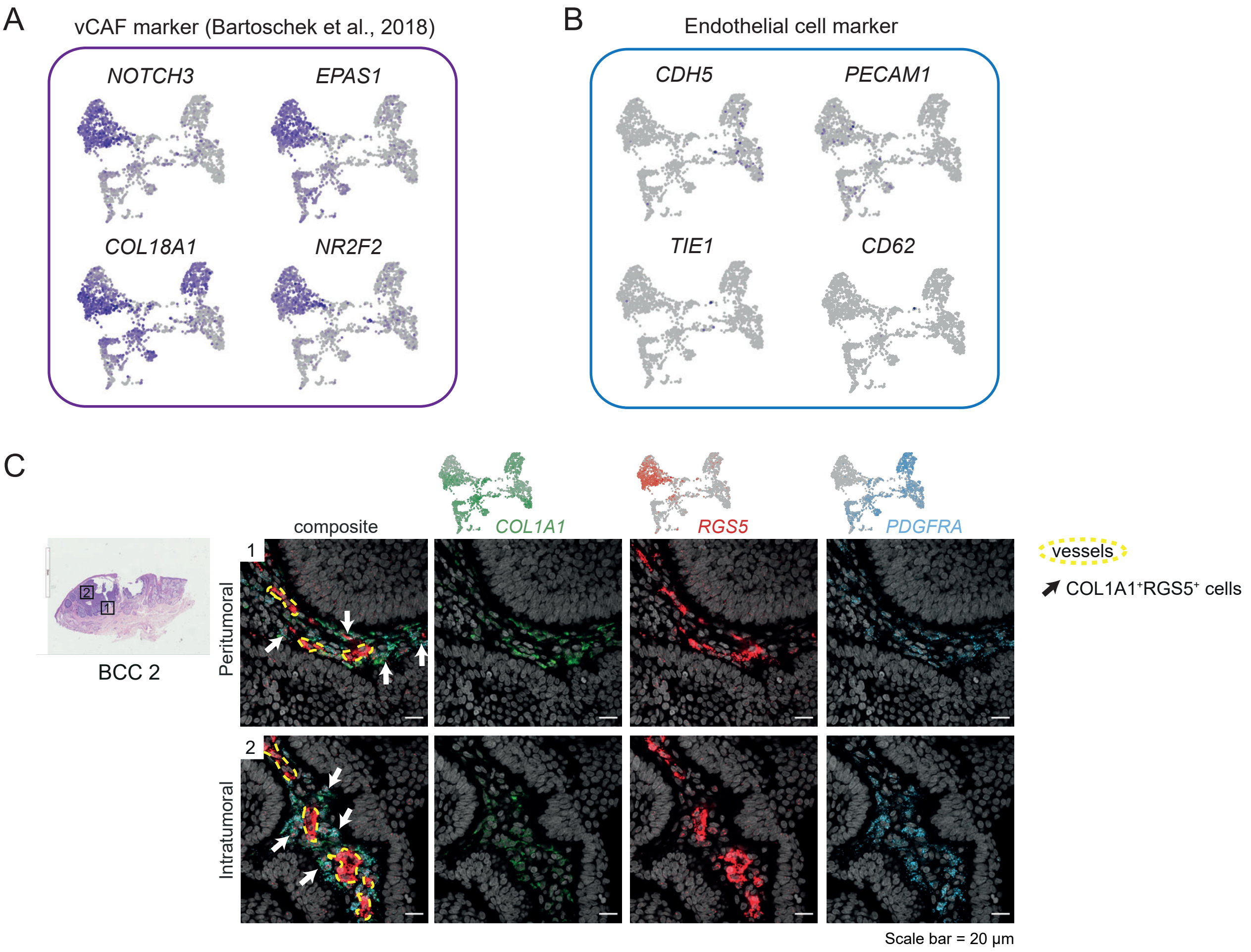

A
vCAF marker (Bartoschek et al., 2018)
NOTCH3
EPAS1
COL18A1
NR2F2
B
Endothelial cell marker
CDH5
PECAM1
TIE1
CD62
C
BCC 2
composite
COL1A1
RGS5
PDGFRA
Peritumoral
Intratumoral
1
2
vessels
COL1A1+RGS5+ cells
Scale bar = 20 µm

Figure S6

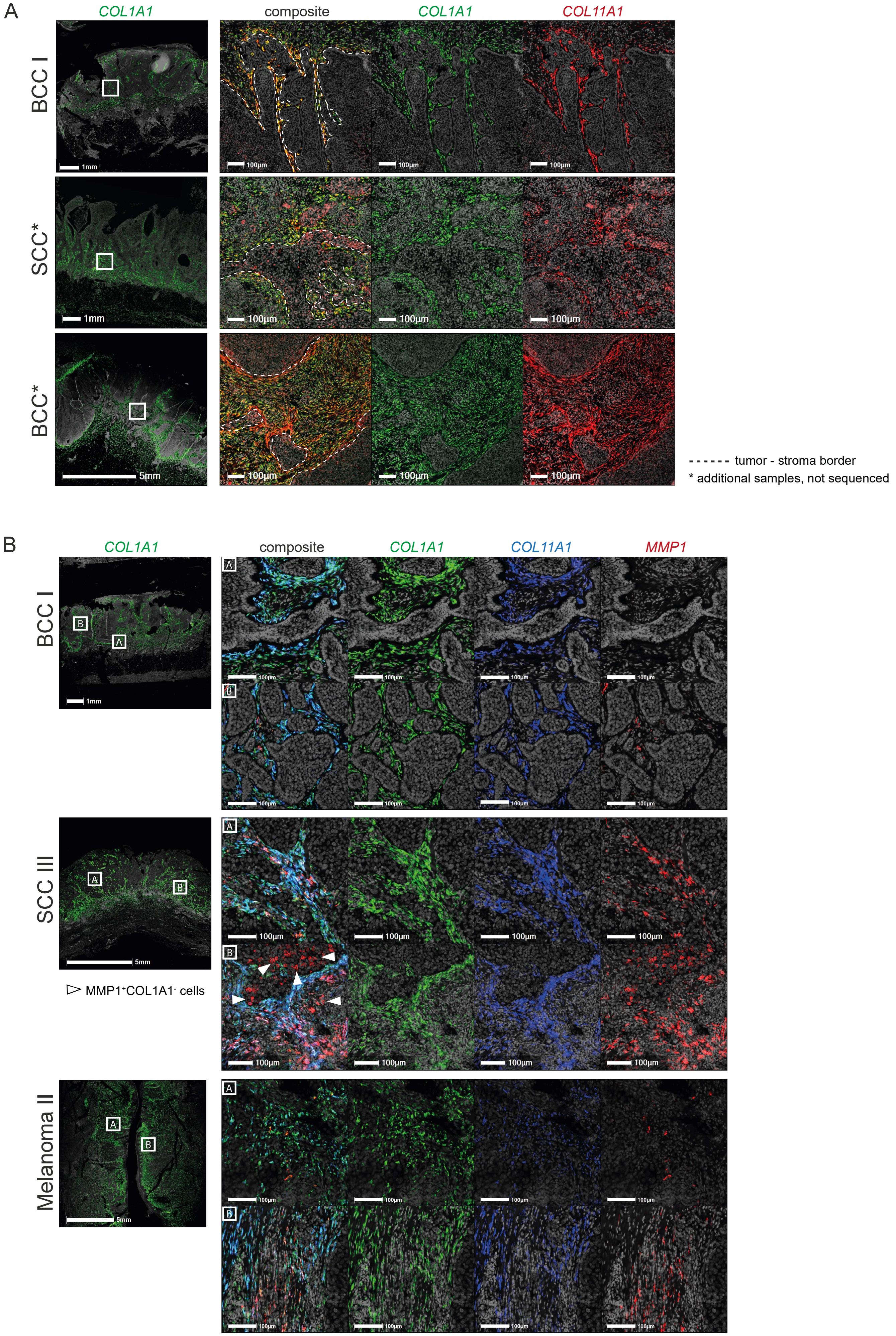
A
COL1A1
composite
COL1A1
COL11A1
BCC I
SCC*
BCC*
1mm
1mm
5mm
100µm
tumor - stroma border
* additional samples, not sequenced
B
COL1A1
composite
COL1A1
COL11A1
MMP1
BCC I
SCC III
Melanoma II
1mm
5mm
5mm
100µm
MMP1+COL1A1- cells

Figure S7

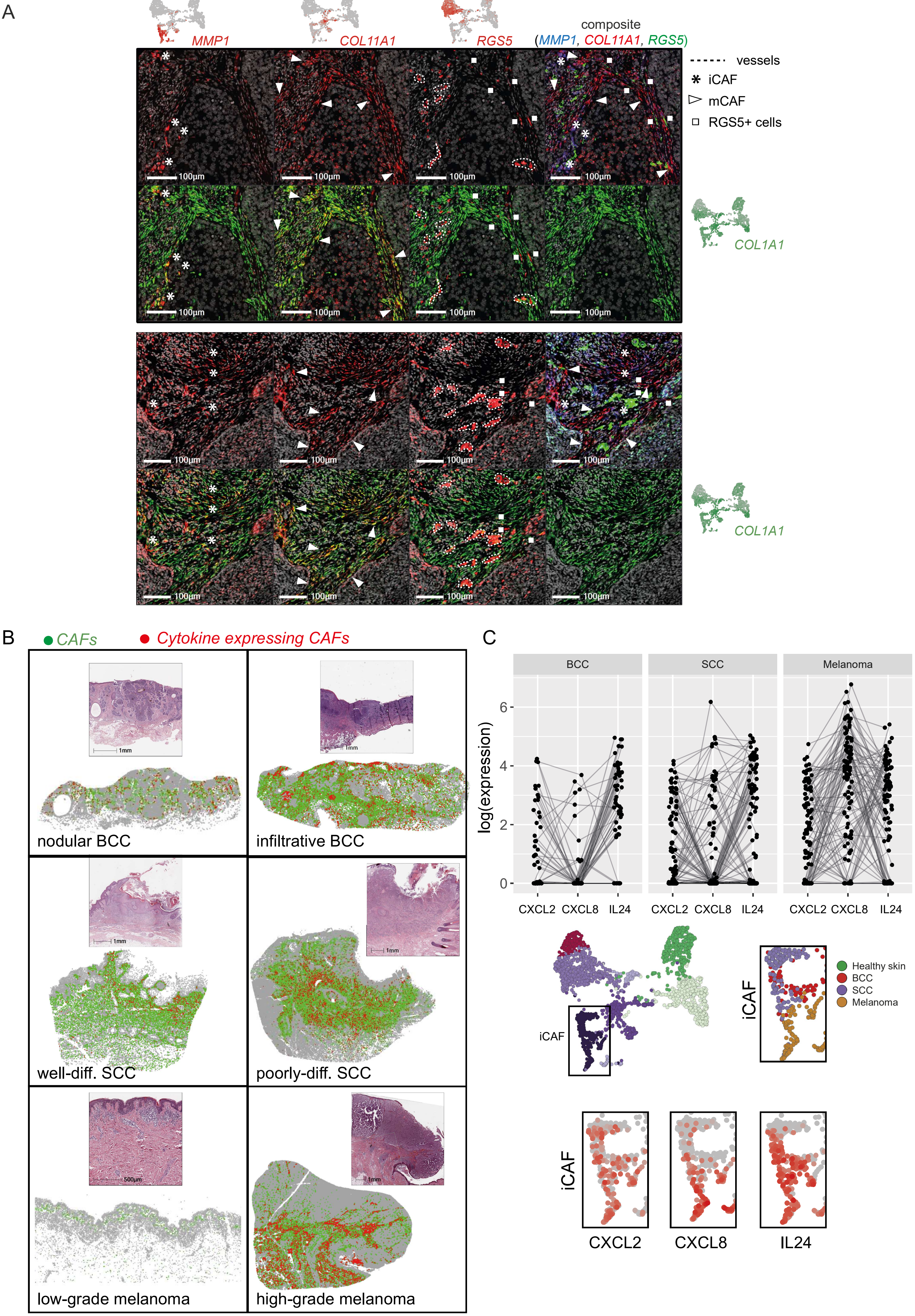
A
MMP1
COL11A1
RGS5
composite
(MMP1, COL11A1, RGS5)
vessels
iCAF
mCAF
RGS5+ cells
100µm
COL1A1
B
CAFs
Cytokine expressing CAFs
1mm
nodular BCC
infiltrative BCC
well-diff. SCC
poorly-diff. SCC
500µm
low-grade melanoma
high-grade melanoma
C
BCC
SCC
Melanoma
log(expression)
6
4
2
0
CXCL2 CXCL8 IL24
iCAF
Healthy skin
BCC
SCC
Melanoma
CXCL2
CXCL8
IL24

Figure S8

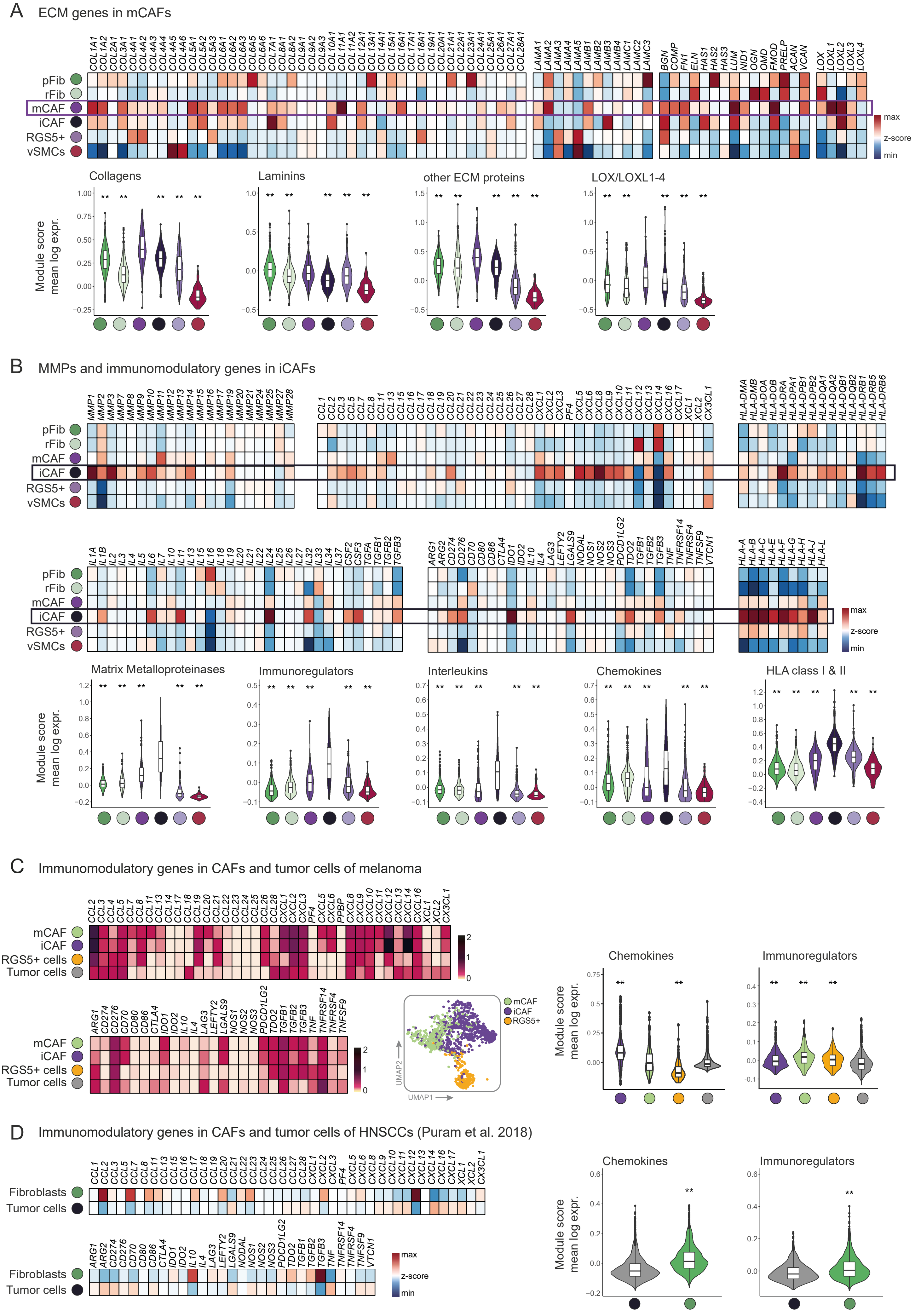
A
ECM genes in mCAFs
COL1A1 COL1A2 COL2A1 COL3A1 COL4A1 COL4A2 COL4A3 COL4A4 COL4A5 COL4A6 COL5A1 COL5A2 COL5A3 COL6A1 COL6A2 COL6A3 COL6A5 COL6A6 COL7A1 COL8A1 COL8A2 COL9A1 COL9A2 COL9A3 COL10A1 COL11A1 COL11A2 COL12A1 COL13A1 COL14A1 COL15A1 COL16A1 COL17A1 COL18A1 COL19A1 COL20A1 COL21A1 COL22A1 COL23A1 COL24A1 COL25A1 COL26A1 COL27A1 COL28A1
LAMA1 LAMA2 LAMA3 LAMA4 LAMA5 LAMB1 LAMB2 LAMB3 LAMB4 LAMC1 LAMC2 LAMC3
BGN COMP FN1 ELN HAS1 HAS2 HAS3 LUM NID1 OGN OMD FMOD PRELP ACAN VCAN
LOX LOXL1 LOXL2 LOXL3 LOXL4
pFib
rFib
mCAF
iCAF
RGS5+
vSMCs
max
z-score
min
Collagens
Laminins
other ECM proteins
LOX/LOXL1-4
Module score
mean log expr.
B
MMPs and immunomodulatory genes in iCAFs
MMP1 MMP2 MMP3 MMP7 MMP8 MMP9 MMP10 MMP11 MMP12 MMP13 MMP14 MMP15 MMP16 MMP17 MMP19 MMP20 MMP21 MMP24 MMP25 MMP27 MMP28
CCL1 CCL2 CCL3 CCL5 CCL7 CCL8 CCL11 CCL13 CCL15 CCL16 CCL17 CCL18 CCL19 CCL20 CCL21 CCL22 CCL23 CCL24 CCL25 CCL26 CCL27 CCL28 CXCL1 CXCL2 CXCL3 PF4 CXCL5 CXCL6 CXCL8 CXCL9 CXCL10 CXCL11 CXCL12 CXCL13 CXCL14 CXCL16 CXCL17 XCL1 XCL2 CX3CL1
HLA-DMA HLA-DMB HLA-DOA HLA-DOB HLA-DRA HLA-DPA1 HLA-DPB1 HLA-DPB2 HLA-DQA1 HLA-DQA2 HLA-DQB1 HLA-DQB2 HLA-DRB1 HLA-DRB5 HLA-DRB6
IL1A IL1B IL2 IL3 IL4 IL5 IL6 IL7 IL10 IL11 IL13 IL15 IL16 IL18 IL19 IL20 IL21 IL22 IL24 IL25 IL26 IL27 IL32 IL33 IL34 IL37 CSF2 CSF3 TGFA TGFB1 TGFB2 TGFB3
ARG1 ARG2 CD274 CD276 CD70 CD80 CD86 CTLA4 IDO1 IDO2 IL10 IL4 LAG3 LEFTY2 LGALS9 NODAL NOS1 NOS2 NOS3 PDCD1LG2 TDO2 TGFB1 TGFB2 TGFB3 TNF TNFRSF14 TNFRSF4 TNFSF9 VTCN1
HLA-A HLA-B HLA-C HLA-E HLA-F HLA-G HLA-H HLA-J HLA-L
Matrix Metalloproteinases
Immunoregulators
Interleukins
Chemokines
HLA class I & II
C
Immunomodulatory genes in CAFs and tumor cells of melanoma
mCAF
iCAF
RGS5+ cells
Tumor cells
UMAP1
UMAP2
RGS5+
Chemokines
Immunoregulators
D
Immunomodulatory genes in CAFs and tumor cells of HNSCCs (Puram et al. 2018)
Fibroblasts
Tumor cells
Chemokines
Immunoregulators

Figure S9

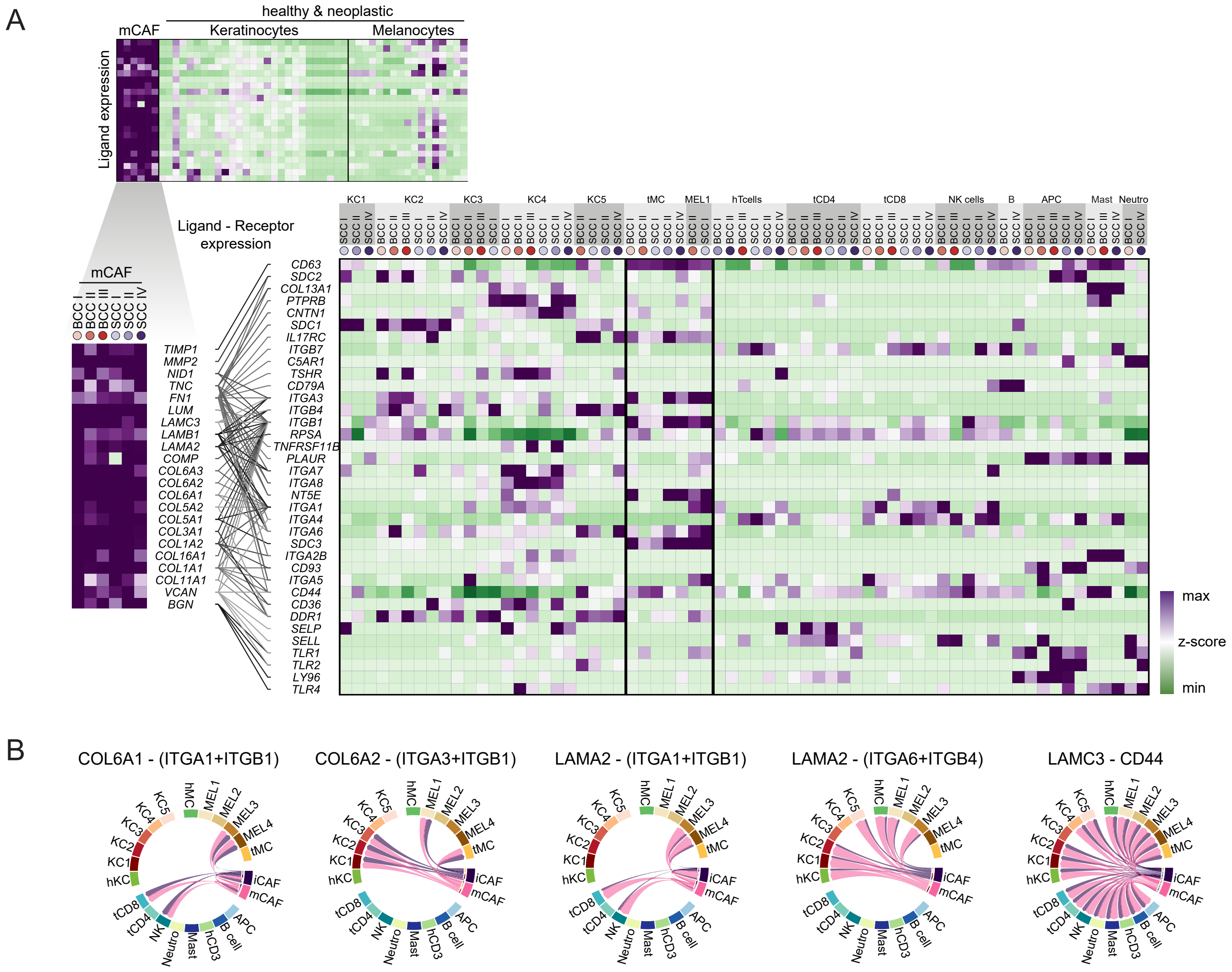
A
healthy & neoplastic
mCAF
Keratinocytes
Melanocytes
Ligand expression
Ligand - Receptor expression
TIMP1
MMP2
NID1
TNC
FN1
LUM
LAMC3
LAMB1
LAMA2
COMP
COL6A3
COL6A2
COL6A1
COL5A2
COL5A1
COL3A1
COL1A2
COL16A1
COL1A1
COL11A1
VCAN
BGN
CD63
SDC2
COL13A1
PTPRB
CNTN1
SDC1
IL17RC
ITGB7
C5AR1
TSHR
CD79A
ITGA3
ITGB4
ITGB1
RPSA
TNFRSF11B
PLAUR
ITGA7
ITGA8
NT5E
ITGA1
ITGA4
ITGA6
SDC3
ITGA2B
CD93
ITGA5
CD44
CD36
DDR1
SELP
SELL
TLR1
TLR2
LY96
TLR4
KC1
KC2
KC3
KC4
KC5
tMC
MEL1
hTcells
tCD4
tCD8
NK cells
B
APC
Mast
Neutro
BCC I
BCC II
BCC III
SCC I
SCC II
SCC IV
max
z-score
min
B
COL6A1 - (ITGA1+ITGB1)
COL6A2 - (ITGA3+ITGB1)
LAMA2 - (ITGA1+ITGB1)
LAMA2 - (ITGA6+ITGB4)
LAMC3 - CD44
hKC
KC1
KC2
KC3
KC4
KC5
hMC
MEL1
MEL2
MEL3
MEL4
tMC
iCAF
mCAF
APC
B cell
hCD3
Mast
Neutro
NK
tCD4
tCD8

Figure S10

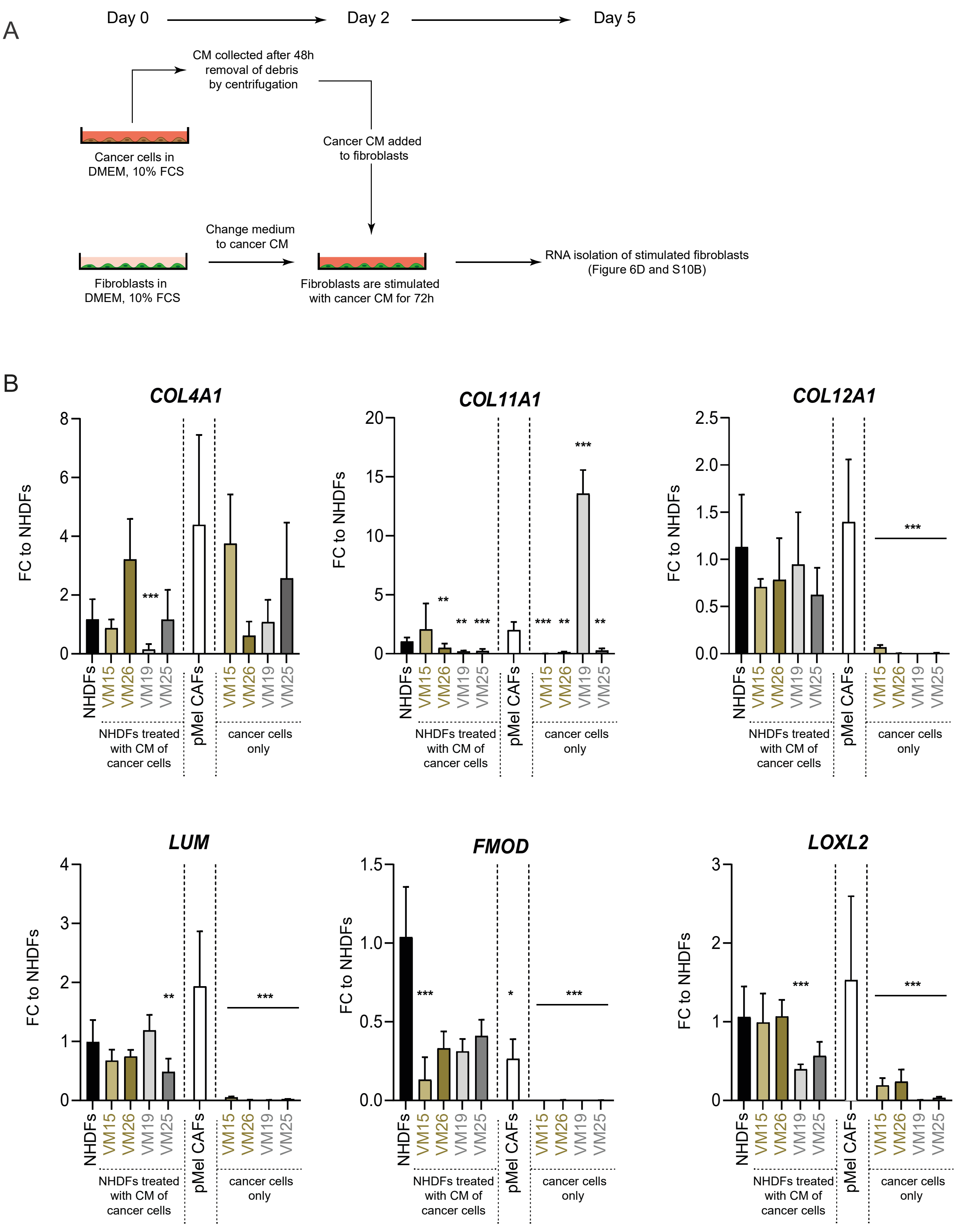


VM08, VM15, VM26: metastatic melanoma cell line
VM19, VM25: primary melanoma cell line
pMel CAFs: fibroblasts isolated from a primary melanoma

Figure S11

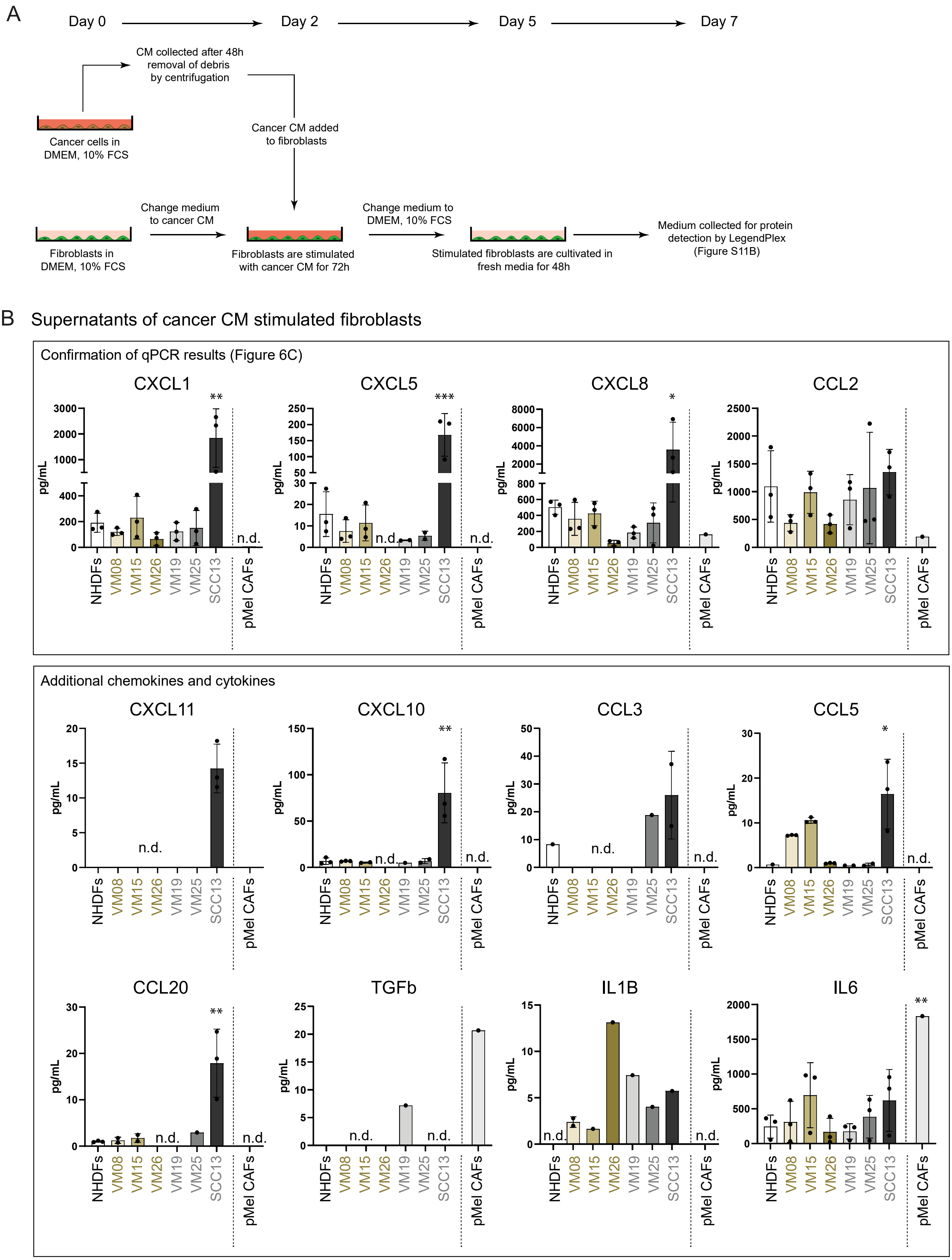


n.d. - not detected/below lowest concentration of standard

### Figure S12

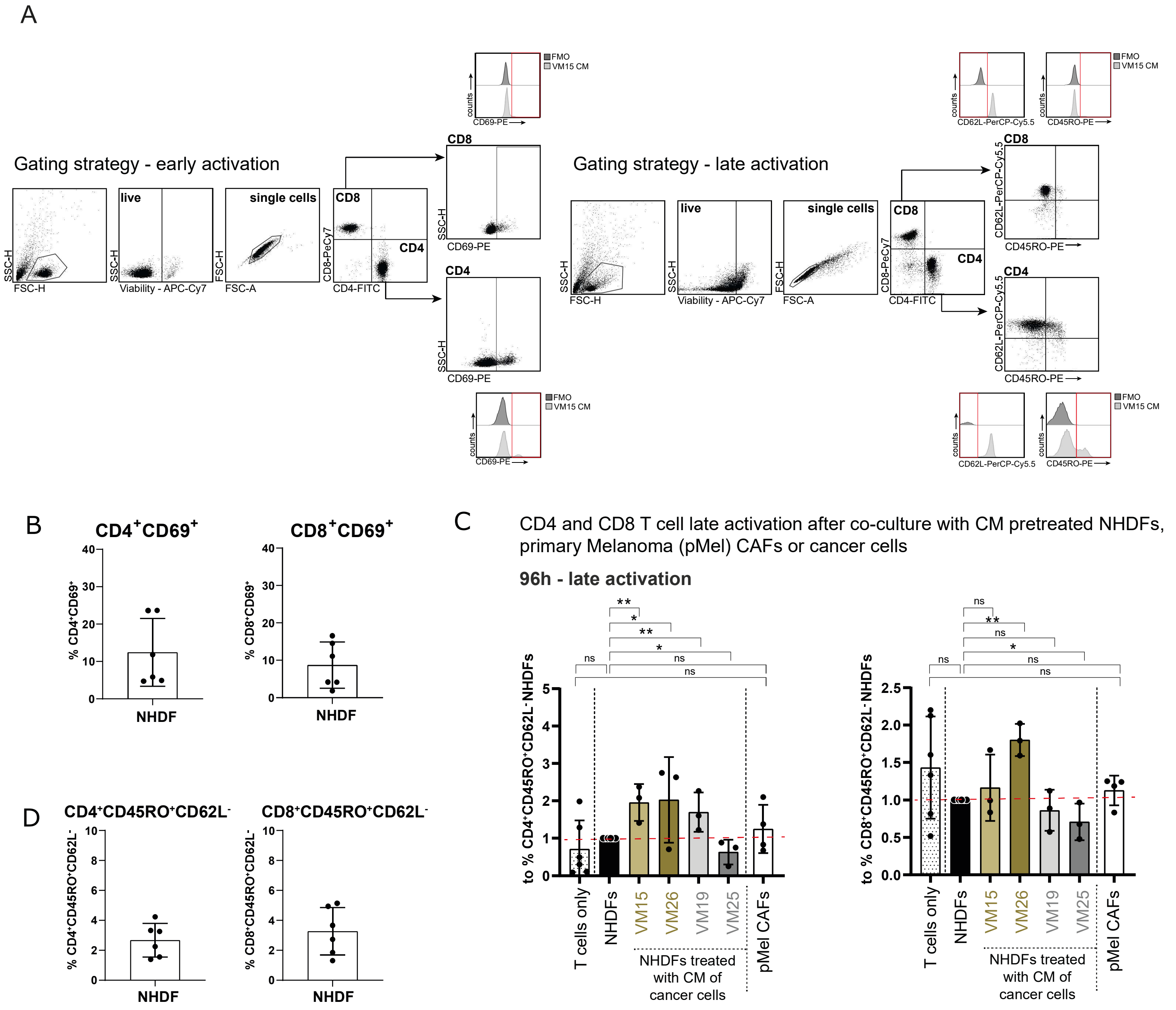
A
Gating strategy - early activation
live
single cells
CD8
CD4
SSC-H
FSC-H
Viability - APC-Cy7
FSC-A
CD8-PeCy7
CD4-FITC
CD69-PE
FMO
VM15 CM
counts
Gating strategy - late activation
CD62L-PerCP-Cy5.5
CD45RO-PE
B
CD4+CD69+
CD8+CD69+
% CD4+CD69+
% CD8+CD69+
NHDF
C
CD4 and CD8 T cell late activation after co-culture with CM pretreated NHDFs, primary Melanoma (pMel) CAFs or cancer cells
96h - late activation
to % CD4+CD45RO+CD62L- NHDFs
to % CD8+CD45RO+CD62L- NHDFs
ns
T cells only
NHDFs
VM15
VM26
VM19
VM25
pMel CAFs
NHDFs treated with CM of cancer cells
D
CD4+CD45RO+CD62L-
CD8+CD45RO+CD62L-
% CD4+CD45RO+CD62L-
% CD8+CD45RO+CD62L-

Figure S13

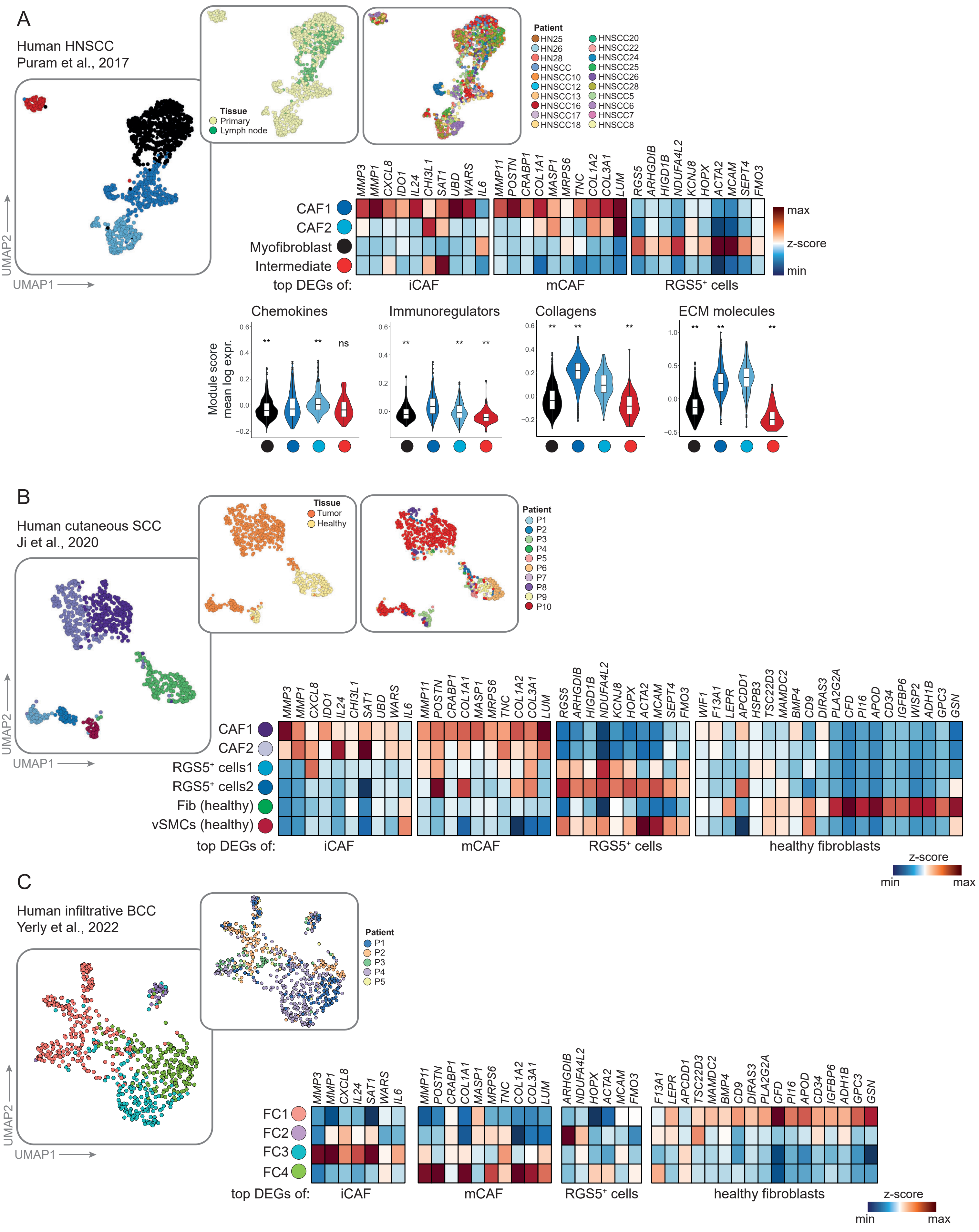

A
Human HNSCC
Puram et al., 2017
Tissue
Primary
Lymph node
Patient
HN25
HN26
HN28
HNSCC
HNSCC10
HNSCC12
HNSCC13
HNSCC16
HNSCC17
HNSCC18
HNSCC20
HNSCC22
HNSCC24
HNSCC25
HNSCC26
HNSCC28
HNSCC5
HNSCC6
HNSCC7
HNSCC8
UMAP2
UMAP1
CAF1
CAF2
Myofibroblast
Intermediate
top DEGs of:
iCAF
mCAF
RGS5+ cells
max
z-score
min
Chemokines
Immunoregulators
Collagens
ECM molecules
Module score
mean log expr.
B
Human cutaneous SCC
Ji et al., 2020
Tissue
Tumor
Healthy
Patient
P1
P2
P3
P4
P5
P6
P7
P8
P9
P10
CAF1
CAF2
RGS5+ cells1
RGS5+ cells2
Fib (healthy)
vSMCs (healthy)
top DEGs of:
iCAF
mCAF
RGS5+ cells
healthy fibroblasts
z-score
min
max
C
Human infiltrative BCC
Yerly et al., 2022
Patient
P1
P2
P3
P4
P5
FC1
FC2
FC3
FC4
top DEGs of:
iCAF
mCAF
RGS5+ cells
healthy fibroblasts
z-score
min
max
